# Cell polarity determinant Dlg1 facilitates epithelial invagination by promoting tissue-scale mechanical coordination

**DOI:** 10.1101/2021.12.14.472598

**Authors:** Melisa A. Fuentes, Bing He

## Abstract

Epithelial folding mediated by apical constriction serves as a fundamental mechanism to convert flat epithelial sheets into multilayered structures. It remains elusive whether additional mechanical inputs are required for folding mediated by apical constriction. Using *Drosophila* mesoderm invagination as a model, we identified an important role for the non-constricting, lateral mesodermal cells adjacent to the constriction domain (“flanking cells”) in facilitating epithelial folding. We found that depletion of the basolateral determinant, Dlg1, disrupts the transition between apical constriction and invagination without affecting the rate of apical constriction. Strikingly, the observed delay in invagination is associated with ineffective apical myosin contractions in the flanking cells that lead to overstretching of their apical domain. The defects in the flanking cells impede ventral-directed movement of the lateral ectoderm, suggesting reduced mechanical coupling between tissues. Specifically disrupting the flanking cells in wildtype embryos by laser ablation or optogenetic depletion of cortical actin is sufficient to delay the apical constriction-to-invagination transition. Our findings indicate that effective mesoderm invagination requires intact flanking cells and suggest a role for tissue-scale mechanical coupling during epithelial folding.

## Introduction

Apical constriction is a commonly used mechanism to achieve epithelial folding during tissue morphogenesis (Martin and Goldstein, 2014; Sawyer et al., 2010). During apical constriction, constricting cell apices shrink and the basal ends of the constricting cells expand, resulting in a characteristic wedge-like shape. Such cell shape changes are accompanied by folding of the epithelial sheet into a 3-dimensional (3D) tissue. Apical constriction mediated epithelial folding occurs in a variety of different developmental contexts, such as neural tube formation in vertebrates (Nishimura et al., 2012), gastrulation in *Drosophila* (Leptin and Grunewald, 1990; Sweeton et al., 1991), endoderm invagination in ascidians (Sherrard et al., 2010), and new branch formation during avian lung morphogenesis (Kim et al., 2013). While we have a detailed understanding of how constriction forces are generated near the apical surface of cells (Martin and Goldstein, 2014; Munjal and Lecuit, 2014), it is less well understood how apical forces drive coordinated cell shape changes in the constricting cells and their non-constricting neighbors to transform a flat epithelium into a multilayered structure.

Folding of the prospective mesoderm during *Drosophila* gastrulation provides an excellent model to study epithelial folding. During gastrulation, a subset of ventrally localized mesodermal cells invaginate from the surface of the embryo to form a furrow (Leptin, 1999). Ventral furrow formation completes within 20 minutes and is characterized by an apical constriction phase and an invagination phase (Sweeton et al., 1991) (Fig. 1A). In *Drosophila* embryos, the presumptive mesoderm is approximately 18 cells wide along the medial-lateral axis. The middle 12 cells that comprise the ventral mesoderm (“constricting mesodermal cells”) undergo apical constriction during gastrulation, whereas the 3 cells flanking each side of the constriction domain that comprise the lateral mesoderm (“flanking cells”) do not. During apical constriction, the constricting mesodermal cells accumulate non-muscle myosin II (henceforth “myosin”) at their apices and elongate in the apical-basal direction (“cell lengthening”). Meanwhile, the constricting mesodermal cells pull the flanking cells towards the ventral midline, causing the flanking cells to become stretched (Sweeton et al., 1991). Shortly thereafter, during invagination, the constricting mesodermal cells undergo shortening and invaginate inwards to form the ventral furrow (“cell shortening”), while the flanking cells remain closer to the surface of the embryo and become the neck of the ventral furrow (Sweeton et al., 1991) (Fig. 1A).

**Figure 1.**
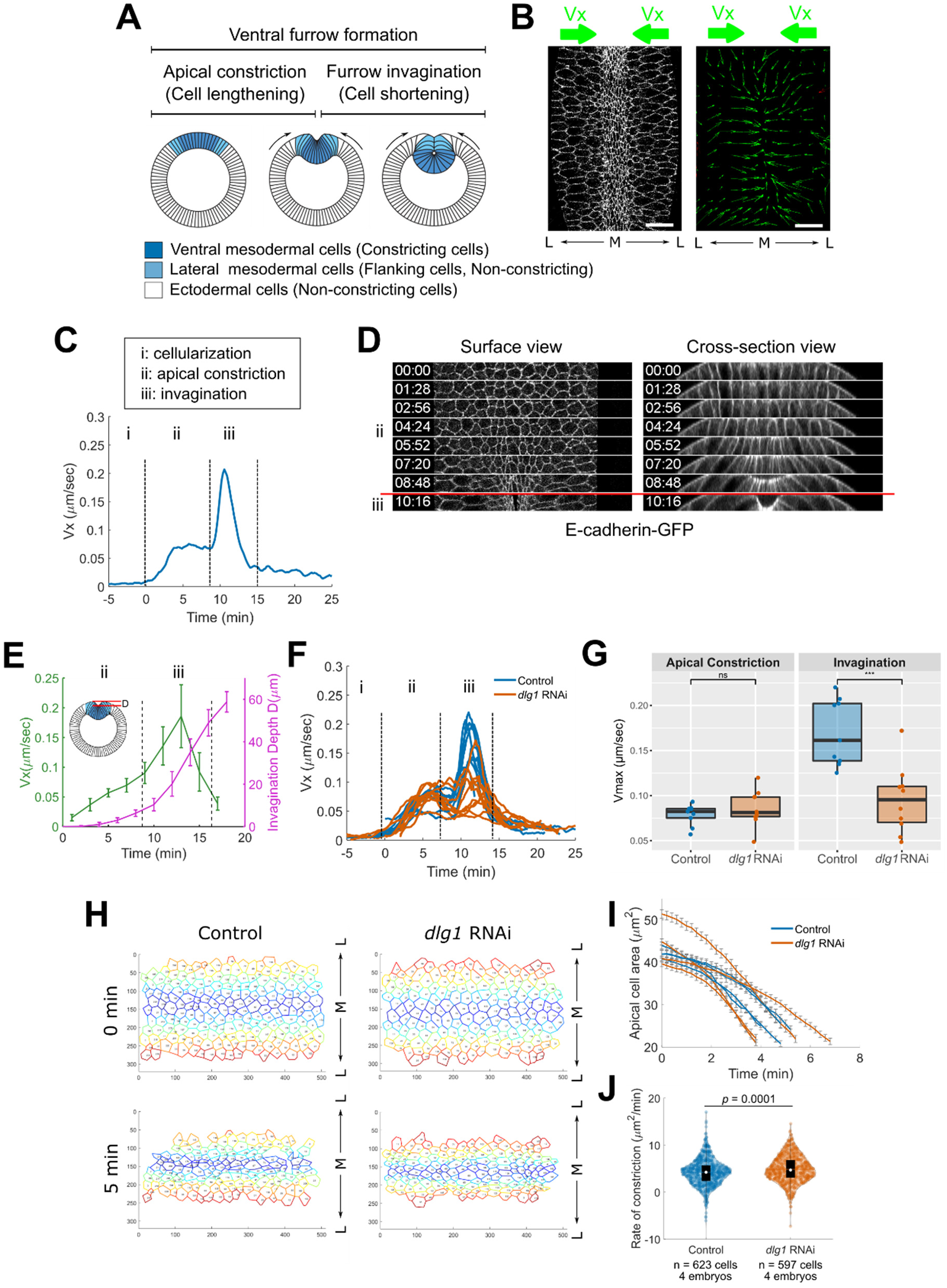
*dlg1* RNAi embryos undergo normal apical constriction but exhibit slower invagination. **(A)** A schematic depicting the behavior of the ventral mesodermal cells (constricting cells; dark blue), the lateral mesodermal cells (flanking cells; light blue), and the ectodermal cells (non-constricting; white) during *Drosophila* ventral furrow formation. Arrows indicate tissue flow towards the ventral midline during mesoderm invagination. **(B)** PIV analysis of tissue movement (green arrows) during ventral furrow formation. A control embryo (left) and its corresponding velocity vectors (right; green arrows) at a single time point are shown as an example. Scale bar: 20 μm. **(C)** The average velocity of tissue movement 10 – 30 μm away from the ventral midline (V_x_; green arrows in B) was plotted over time. Phases i, ii, and iii correspond to cellularization, apical constriction, and invagination, respectively. Time 0 marks the onset of gastrulation throughout the text unless otherwise noted. **(D)** Movie stills showing the surface view and cross-section view of a control embryo undergoing gastrulation. Ventral side is facing up. T = 0:00 (mm:ss) is the onset of gastrulation. Red line demarcates the transition from apical constriction to invagination. **(E)** Measurement of the velocity of tissue movement 30 μm away from the ventral midline (V_x_) and invagination depth, D, in the control embryos (n = 8) imaged on a multiphoton microscope. **(F-G)** Maximal V_x_ during apical constriction is comparable between control (n = 9) and *dlg1* RNAi embryos (n = 10), whereas maximal V_x_ during invagination is substantially reduced in the *dlg1* RNAi embryos. Student t-test, unpaired, two-tailed. *: p <= 0.05; **: p <= 0.01; ***: p <= 0.001; ****: p <= 0.0001. **(H)** Individual cells within the constriction domain were segmented in order to measure cell area reduction over time during apical constriction. The constriction cell domain at T = 0 and T = 5 minutes is shown. **(I)** Quantification of average cell area reduction in the constriction domain over time in control (n = 4) and *dlg1* RNAi embryos (n = 4). Each curve represents one embryo (n = 129 to 164 cells per embryo). Error bar: standard error of the mean. **(J)** Distribution of apical constriction rate for individual cells in control and *dlg1* RNAi embryos. Cells from embryos with the same genotype were pooled.

The molecular mechanism that regulates apical accumulation of myosin, and thus apical constriction, in the constricting mesodermal cells has been extensively studied. The expression of two transcription factors, Twist and Snail, at the ventral side of the embryo specifies mesodermal cell fate and promotes ventral furrow formation (Leptin, 1991). Twist and Snail activate the recruitment of RhoGEF2, a Rho GTPase activator, to the apex of the mesodermal precursor cells via a G-protein coupled receptor pathway (Costa et al., 1994; Kerridge et al., 2016; Kölsch et al., 2007; Manning et al., 2013; Parks and Wieschaus, 1991). RhoGEF2 then activates myosin across the apical surface of the prospective mesoderm through the Rho-Rho kinase pathway (Barrett et al., 1997; Dawes-Hoang et al., 2005; Häcker and Perrimon, 1998; Martin et al., 2009; Mason et al., 2013; Nikolaidou and Barrett, 2004). Upon activation, myosin forms a contractile actomyosin network that undergoes stochastic, pulsatile contractions, which power constriction of cell apices (Martin and Goldstein, 2014; Martin et al., 2009). Myosin activation occurs in a gradient, which contributes to the differential cell behavior observed in the ventral and lateral mesoderm. The constricting mesodermal cells and the flanking cells are characterized by high and low levels of Fog signaling, respectively (Costa et al., 1994). Inhibition of Fog signaling in the flanking cells is mediated by the GPCR kinase, Gprk2 (Fuse et al., 2013).

A number of recent studies have shown that in addition to apical myosin, myosin localized at other subcellular locations in the constricting cells can also influence invagination of the ventral furrow. It has been proposed, and recently demonstrated experimentally, that accumulation of myosin along the lateral cortex of the constricting cells facilitates cell shortening and furrow invagination by increasing tension along the apical-basal axis (Gracia et al., 2019; John and Rauzi, 2021). In addition, it has been proposed that proper downregulation of myosin at the basal cortex of the mesodermal cells facilitates tissue internalization by softening the basal membrane (Polyakov et al., 2014). In support of this view, a recent study has shown that prolonged retention of basal myosin in the mesodermal cells inhibits ventral furrow invagination (Krueger et al., 2018). Thus, appropriate regulation of myosin contractility in the constricting cells is essential for apical constriction and invagination. Up to this point, it remains unknown whether mechanical factors other than myosin contractility in the constricting cells also contribute to invagination of the ventral furrow.

Interestingly, a few recent studies have reported that perturbation of the non-constricting tissues adjacent to the constriction domain negatively impacts ventral furrow invagination. One study found that increasing contractility in the flanking cells by mutating Gprk2 causes the flanking cells to undergo apical constriction instead of stretching (Fuse et al., 2013). Constriction of the flanking cells hinders ventral furrow invagination and results in an open, U-shaped furrow (Fuse et al., 2013). It has also been shown that ectopically upregulating apical contractility in the neighboring lateral ectodermal cells reduces the speed of ventral furrow invagination and causes the ventral furrow to unfold (Perez-Mockus et al., 2017). Furthermore, when movement of the neighboring ectodermal tissues is inhibited by cauterization, ventral furrow invagination is obstructed (Rauzi et al., 2015). Together, these studies demonstrate that mesoderm invagination can be influenced by the mechanical properties of the surrounding non-constricting tissues. However, the exact mechanical contribution of the non-constricting tissues during ventral furrow formation is currently unknown.

In this work, we used a quantitative live-imaging based assay to search for mutants that undergo normal apical constriction, but have defects in tissue invagination during ventral furrow formation. We found that depletion of Dlg1, a protein important for establishing and maintaining apical-basal polarity in epithelial cells, resulted in the desired phenotype. In several types of mature epithelia in *Drosophila*, Dlg1 helps maintain polarity and basolateral identity by promoting septate and adherens junction formation and by preventing basal expansion of apical polarity proteins (Tanentzapf and Tepass, 2003; Woods et al., 1996). In addition, during polarity establishment in the *Drosophila* blastoderm, Dlg1 facilitates the recruitment of apical polarity determinants to the adherens junctions and promotes adherens junction assembly (Bonello et al., 2019). We found that RNAi-mediated knockdown of Dlg1 impairs invagination of the ventral furrow, without noticeably affecting myosin contractility in the constricting cells or their morphology. In *dlg1* RNAi embryos, the flanking cells become hyper-stretched as the ventral mesodermal cells constrict apically. The altered behavior of the flanking cells is associated with a reduced level of apical adherens junctions and ineffective apical myosin contractions that fail to restrain apical area when the flanking cells are pulled on by the constricting cells. At the tissue level, hyper-stretching of the flanking cells weakens coupling between apical constriction and ventral-oriented movement of the neighboring ectodermal tissue. This impaired coupling is associated with a delay in the transition between apical constriction and invagination during ventral furrow formation. In wildtype embryos, disruption of the non-constricting cells adjacent to the constriction domain by targeted laser ablation or optogenetic downregulation of the cortical actomyosin network results in a similar delay in invagination as observed in the *dlg1* RNAi embryos. Together, these results demonstrate the importance of the flanking cells during ventral furrow formation and suggest that mechanical coupling between the constriction domain and the surrounding ectodermal tissue promotes robust furrow invagination. Furthermore, our findings reveal a previously unappreciated function for apical myosin contractions in stabilizing flanking cell apical morphology, which depends on the function of Dlg1.

## Results

### Establishing a quantitative live-imaging approach to monitor tissue flow during Drosophila ventral furrow formation

Inspired by the observation that apical constriction and invagination are temporally distinct processes during ventral furrow formation (Polyakov et al., 2014; Rauzi et al., 2015), we hypothesized that invagination may have its own unique regulatory inputs separate from those of apical constriction. To search for mutants that are specifically defective in furrow invagination but not in apical constriction, we developed a live imaging-based approach that allowed us to monitor tissue movement at the surface of the embryo during ventral furrow formation. We focused on a relatively shallow region near the surface of the ventral tissue so that we could analyze apical cell dynamics with relatively high spatial and temporal resolution. To visualize cell membranes, we imaged embryos expressing the GFP-tagged adherens junction marker, E-cadherin (E-cadherin-GFP). The embryos also expressed mCherry-tagged myosin regulatory light chain, Spaghetti Squash (Sqh-mCherry), which allowed us to examine myosin dynamics. To account for the curvature of the embryo, a “flattened” surface projection was generated for each embryo post-acquisition, which allowed us to easily visualize tissue flow from the lateral regions towards the ventral midline (Heemskerk and Streichan, 2015).

Following generation of the surface view, we used a particle image velocimetry (PIV) software (Taylor et al., 2010) to track the movement of the tissue in a region spanning 10 ‒ 30 μm from the ventral midline during gastrulation (Fig. 1A-B; green arrows). We focused on tissue velocity along the medial-lateral axis (“V_x_”) because it is the predominant direction of tissue movement during ventral furrow formation. As expected, we found that there is little cell movement during cellularization (V_x_ = 0; Fig. 1C; part i). After the onset of apical constriction (which we defined as T = 0 minutes for the remainder of the text, unless otherwise noted), constriction of the ventral mesodermal cells initiates tissue movement towards the ventral midline, resulting in an increase in V_x_ that peaks/plateaus approximately 5 minutes into gastrulation (Fig. 1C; part ii). Approximately 8 minutes after gastrulation onset, V_x_ showed another rapid increase, forming a second, more prominent peak between T = 8 minutes and T = 15 minutes (Fig. 1C; part iii). The rapid increase in V_x_ shown in part iii of the curve temporally correlates with the onset of rapid invagination (Fig. 1D; red line). To further confirm the relationship between V_x_ and furrow invagination, we acquired movies with a multiphoton microscope, which allowed us to image the entire ventral furrow (up to 100 μm deep) at a lower temporal resolution. Quantification of V_x_ and invagination depth (“D”) over time confirmed the rapid increase in V_x_ during the invagination phase (Fig. 1E). Thus, the velocity of tissue flow towards the ventral midline provides a sensitive readout for the progress of ventral furrow formation and reveals the rate of tissue flow during different phases of furrow formation.

### Knockdown of Dlg1, a basolateral polarity determinant, impairs the invagination phase of ventral furrow formation without causing obvious defects in apical constriction

Using the approach mentioned above, we examined embryos containing maternally-loaded short hairpin RNAs to silence the expression of specific candidate genes important for epithelial polarity and cell-cell adhesion (Baz, aPKC, Par-6, Crb, Sdt, Dlg, Scrib, Lgl, Shg, Vang) (Ni et al., 2011). In this small-scale screen, we searched for mutants which exhibited normal tissue flow during apical constriction, but significantly reduced tissue flow during invagination. We found that knockdown of the basolateral determinants, Dlg1, Scrib, and Lgl, resulted in the desired phenotype (Fig. 1F; Fig. S1; Movie 1). Since the basolateral determinants often have similar mutant phenotypes (Bilder et al., 2000), we focused on Dlg1 for the subsequent analysis. Knockdown of Dlg1 was confirmed by immunostaining with an anti-Dlg1 antibody (Fig. S2). The majority of *dlg1* RNAi embryos exhibited normal cellularization, a normal peak of V_x_ in the apical constriction phase, and a reduced peak of V_x_ in the invagination phase (Fig. 1F). We noticed that a small fraction of the embryos showed more severe phenotypes, ranging from abnormal apical constriction to impaired cellularization (∼ 22%, 8 of 37 embryos imaged). The variation in the severity of the phenotype might be attributed to variation in Dlg1 knockdown levels between embryos. Since our goal was to investigate the mechanism of invagination, we focused our analysis on *dlg1* RNAi embryos that showed no visible defects during cellularization and apical constriction. Further quantification confirmed that this group of *dlg1* RNAi embryos had a similar average peak value of V_x_ during the apical constriction phase as control embryos (Fig. 1G). However, the average peak value of V_x_ during the invagination phase in the *dlg1* RNAi embryos is significantly lower than in the control embryos (Fig. 1G). To further confirm that apical constriction is normal in these *dlg1* RNAi embryos, we segmented the apical cell outline of the constricting cells and tracked cell shape changes during apical constriction using Embryo Development Geometry Explorer (EDGE), a MATLAB based image segmentation tool (Methods) (Gelbart et al., 2012). This analysis revealed that the average rate of apical cell area reduction during apical constriction is largely comparable between the control and *dlg1* RNAi embryos (Fig. 1H-I; n = 4 embryos for each genotype).

### Knockdown of Dlg1 results in defects in polarity establishment during cellularization

We performed additional experiments to test whether *dlg1* RNAi embryos show expected polarity defects in early embryos. In the *dlg1* RNAi embryos, several proteins that are normally localized subapically at the end of cellularization, including Bazooka/Par-3 (Baz), Canoe/Afadin (Cno), and E-cadherin, exhibited a noticeable basal shift along the lateral membrane (Fig. S3). These observations are consistent with the recently reported role of Dlg1 in facilitating the apical enrichment of apical polarity determinants during cellularization (Bonello et al., 2019). In addition, we found that similar to other subapical polarity markers, F-actin as visualized by the Venus-tagged actin binding domain of Utrophin (Utr-Venus), also became more basally shifted in *dlg1* RNAi embryos compared to control embryos at the end of cellularization (Fig. S4A-D). The defect in F-actin distribution at this stage was not observed in Bonello et al., perhaps due to differences in the methods used to visualize F-actin (Bonello et al., 2019). To further test the role of Dlg1 in actin localization, we examined Utr-Venus in embryos derived from *dlg1^2^/dlg1^5^* trans-heterozygous mutant females. *dlg1^2^* is a temperature sensitive allele that shows progressively worse phenotypes from 18°C to 29°C, whereas *dlg1^5^* is a hypomorphic allele that is weakly temperature sensitive (Perrimon, 1988). We found that Utr-Venus is more basally spread in *dlg1^2^/dlg1^5^* embryos at 22°C (Fig. S4E-G), consistent with our observations in *dlg1* RNAi embryos. These results suggest that polarized localization of F-actin is sensitive to perturbations in Dlg1 function. We also noticed that the basal spread of Utr-Venus is more severe in the *dlg1^2^/dlg1^5^* embryos than in the *dlg1* RNAi embryos (Fig. S4), suggesting that the *dlg1* RNAi lines used in our analysis act like relatively mild *dlg1* alleles. Together, our analyses indicate that knockdown of Dlg1 generates expected polarity phenotypes in early embryos.

### dlg1 RNAi embryos exhibit a delay in the transition between apical constriction and invagination

To further characterize the invagination phenotype in the *dlg1* RNAi embryos, we used multiphoton-based deep-tissue imaging to visualize the entire depth of the ventral furrow over the course of gastrulation (Fig. 2A; Movie 2). Consistent with our previous observations, in most *dlg1* RNAi embryos the apical constriction rate was similar to that of the control embryos (Fig. 2B; ventral cells achieved 70% apical area reduction in 8.3 ± 1.8 minutes in *dlg1* RNAi embryos, n = 8 embryos, and 7.4 ± 1.5 minutes in control embryos, n = 4 embryos, mean ± s.d.). In control embryos, the constricting mesodermal cells form a shallow furrow with a cup-shaped apical indentation at about 8 – 10 minutes after the onset of apical constriction. Immediately after this “transition” timepoint (“T_trans_”), the tissue undergoes rapid invagination until the furrow is fully internalized (Fig. 2A). In *dlg1* RNAi embryos, the apex of the shallow furrow remains close to the embryo’s surface for a longer period of time before it enters the rapid invagination phase (Fig. 2A; magenta boxes). To further quantify this phenotype, we measured the invagination depth, D, over time, which is defined as the distance between the apex of the furrow and the surface of the embryo (Fig. 2C). In *dlg1* RNAi embryos, it takes a significantly longer time for the ventral furrow to reach an invagination depth of 30 μm compared to control embryos (Fig. 2D; 15.4 ± 3.4 minutes in *dlg1* RNAi (line 1) embryos, n = 20 embryos, 17.1 ± 3 minutes in *dlg1* RNAi (line 2) embryos, n = 9 embryos, and 12.6 ± 1 minutes in control embryos, n = 21 embryos, mean ± s.d.). In addition, the ventral furrow invaginates less deeply in the *dlg1* RNAi embryos, although the final morphology of the ventral furrow is largely normal (Fig. 2E; 53 ± 4 μm in *dlg1* RNAi (line 1) embryos, n = 20 embryos, 52 ± 6 μm in *dlg1* RNAi (line 2) embryos, n = 9 embryos, and 65 ± 5 μm in control embryos, n = 21 embryos, mean ± s.d.). Interestingly, there is a moderate anti-correlation between the delay in invagination and the final invagination depth (Fig. 2F), suggesting that the two phenotypes are linked.

**Figure 2.**
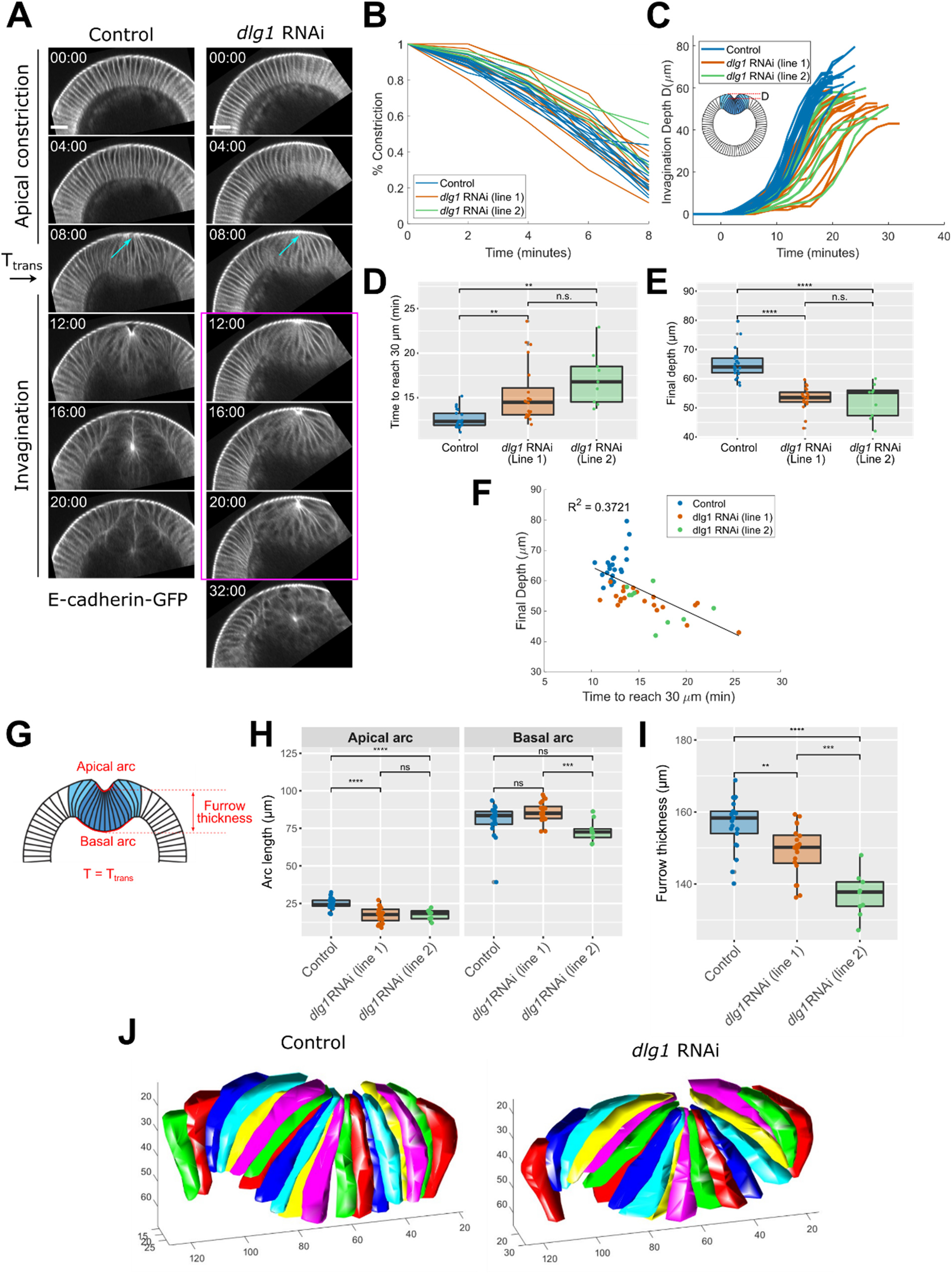
*dlg1* RNAi embryos exhibit a delay in the transition between apical constriction and invagination, and invaginate less deeply. (**A**) Cross-section views of a control and *dlg1* RNAi mutant embryo during ventral furrow formation. E-cadherin-GFP was used as a membrane marker. T = 0:00 (mm:ss) is the onset of gastrulation. Scale bars: 20 μm. Although apical constriction appears normal in the *dlg1* RNAi embryo (cyan arrows), there is a delay in the transition from apical constriction to invagination (magenta box). (**B**) Quantification of the normalized constriction cell domain width over time in the control (n = 9) and *dlg1* RNAi embryos (n = 11). (**C**) Quantification of the invagination depth, D, of the ventral furrow (defined as the distance between the eggshell and the furrow apex) over time in the control (n = 21) and in two *dlg1* RNAi lines (n = 20 & 9). (**D**) The time it takes for the ventral furrow to invaginate 30 µm, T_30µm_, in the control and in the two *dlg1* RNAi lines shown in C. Student t-test, unpaired, two-tailed. *: p <= 0.05; **: p <= 0.01; ***: p <= 0.001; ****: p <= 0.0001. (**E**) Final invagination depth in the control (n = 21) and in the two *dlg1* RNAi lines (n = 20 & 9) shown in C. Student t-test, unpaired, two-tailed. *: p <= 0.05; **: p <= 0.01; ***: p <= 0.001; ****: p <= 0.0001. (**F**) The final invagination depth and T_30µm_ show an anti-correlation. (**G**) Cartoon depicting the measurement of apical and basal arc lengths (outlined in red), as well as furrow thickness at T_trans_ (8 min). Arc lengths were obtained by outlining the ventral furrow from the cross-section view in embryos expressing E-cadherin-GFP. Furrow thickness is defined as the distance between the apex of the ventral furrow and the base of the ventral furrow. (**H, I**) Apical and basal arc length and furrow thickness comparisons for the control (n = 21) and two *dlg1* RNAi lines (n = 20 & 9) at T_trans_. Student t-test, unpaired, two-tailed. *: p <= 0.05; **: p <= 0.01; ***: p <= 0.001; ****: p <= 0.0001. (**J**) 3D segmentation of the constricting cells and their non-constricting neighbors at T_trans_.

Next, we asked whether the constricting, ventral mesodermal cells display any morphological defects prior to T_trans_. First, we measured the length of the curved apical and basal surfaces (“arc length”) of the intermediate furrow at T_trans_ from the cross-section view of the embryo (Fig. 2G). The apical arc length was on average 20% shorter in the *dlg1* RNAi embryos compared to the control embryos, whereas the basal arc length was similar between the two genotypes (Fig. 2H). Next, we estimated the apical-basal thickness of the intermediate furrow at T_trans_ by measuring the height of the constricting cells located at the ventral midline (Fig. 2G). We found that the intermediate furrow in the *dlg1* RNAi embryos was on average 6% thinner than in the control embryos (Fig. 2I). Finally, we performed 3D segmentation and reconstruction of the constricting cells at T_trans_. Our data show that other than the mild reduction in apical-basal cell length, the morphology of the constricting cells and their spatial arrangement in the intermediate furrow in the *dlg1* RNAi embryos is largely comparable to that of the control embryos (Fig. 2J). Together, these data indicate that ventral furrow formation in the *dlg1* RNAi embryos is relatively normal during the apical constriction phase. However, the transition between apical constriction and invagination is impaired in the *dlg1* RNAi embryos, resulting in a delay in furrow invagination.

It is worth noting that the expression of the invagination phenotype upon Dlg1 knockdown is sensitive to the GAL4 driver line that we used to drive the expression of *dlg1* shRNA. The driver line used in our original screen, which contained two copies of *GAL4*, one copy of *E-cadherin-GFP*, and one copy of *Sqh-mCherry*, produced the most prominent invagination phenotype when combined with UAS *dlg1* shRNA. We used this driver line for all subsequent analysis, unless otherwise noted. It is unclear whether this phenomenon is associated with the expression level of GAL4 or the fluorescent markers specific to this driver. Nevertheless, the observed phenotype provided a useful starting point for us to further investigate the factors that are important for an effective transition between apical constriction and invagination during ventral furrow formation.

### The distribution of cortical myosin in the constricting cells is largely normally in the dlg1 RNAi embryos during ventral furrow formation

Although the ventral mesodermal cells undergo relatively normal apical constriction in the *dlg1* RNAi embryos, we wondered whether other defects within these cells could contribute to the invagination defects. In particular, recent studies have shown that in addition to apical myosin, two other subpopulations of myosin in the ventral mesodermal cells can also influence ventral furrow invagination (Fig. 3A). Specifically, accumulation of lateral myosin in the constricting cells bolsters tissue bending by increasing tension along the apical-basal direction, and downregulation of basal myosin in the constricting cells enables tissue bending by decreasing tissue stiffness at the base of the furrow (Gracia et al., 2019; John and Rauzi, 2021; Krueger et al., 2018; Polyakov et al., 2014). We therefore asked whether knockdown of Dlg1 affects lateral or basal myosin in the constricting cells. First, we examined Sqh-mCherry signal at the basal surface of the tissue. Constricting cells in the *dlg1* RNAi embryos underwent a similar basal myosin loss as in the control embryos (Fig. 3B; yellow arrowheads). Next, we examined Sqh-mCherry signal along the lateral membrane of the constricting mesodermal cells. In both the control and *dlg1* RNAi embryos, lateral Sqh-mCherry was sparse, but detectable in the constricting cells soon after the onset of gastrulation (Fig. 3C; red arrows). We did not detect any notable differences in the abundance of lateral myosin between the control and the *dlg1* RNAi embryos. In addition to basal and lateral myosin, we also examined the accumulation of apical myosin in the ventral mesodermal cells. In accordance with our observation that the rate of apical constriction is relatively normal in the *dlg1* RNAi embryos, we found that apical myosin accumulation and distribution is similar between *dlg1* RNAi and control embryos (Fig. 3D-E; Movie 3). Furthermore, we did not detect an obvious increase in local breaks within the supracellular myosin network in the *dlg1* RNAi embryos, suggesting that the network is properly interconnected (Movie 3). Taken together, these observations suggest that the invagination delay in the *dlg1* RNAi embryos is not caused by defects in the activation or spatial organization of myosin in the constricting mesodermal cells.

**Figure 3.**
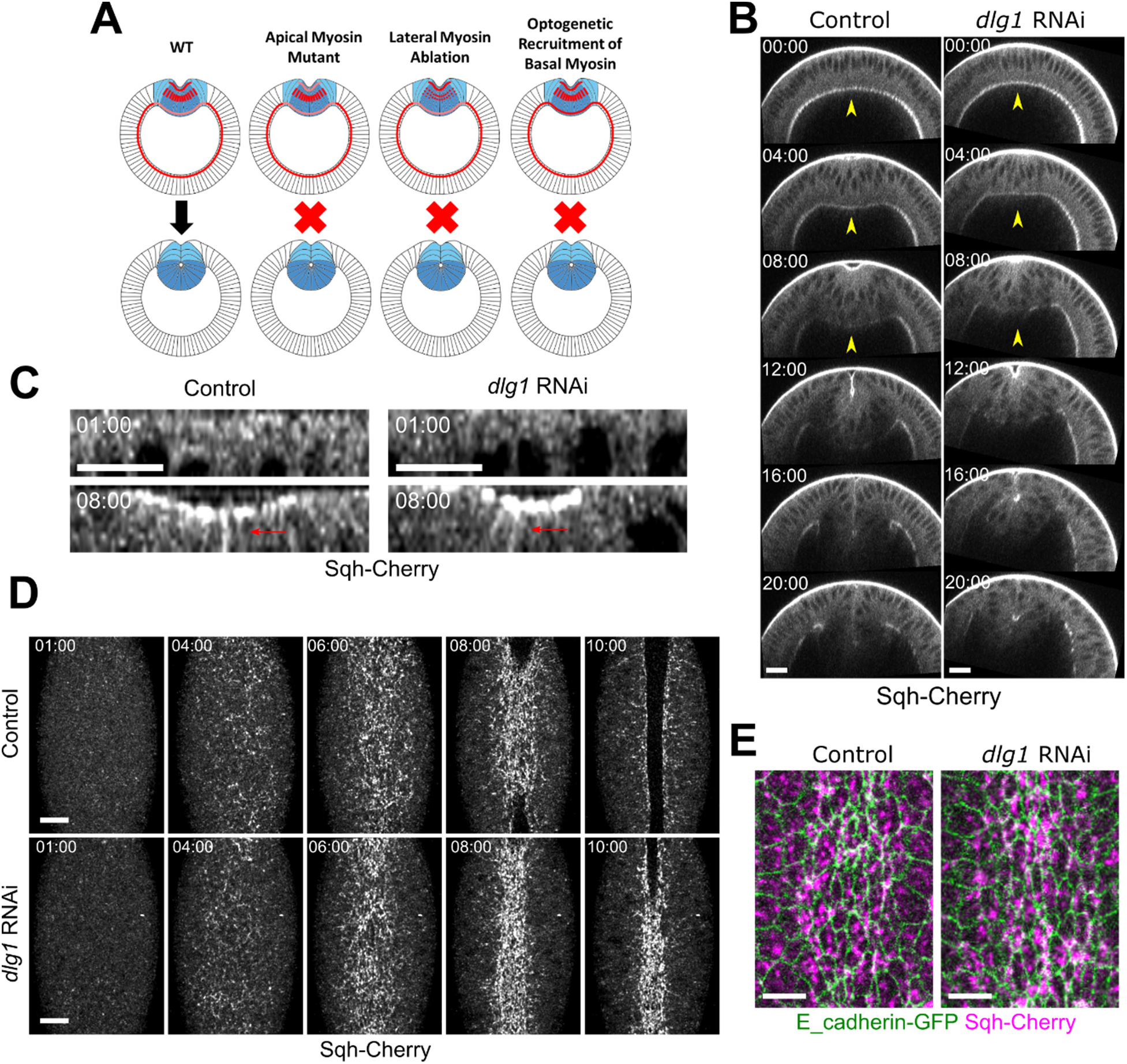
Invagination defects in *dlg1* RNAi embryos are not likely due to abnormal myosin distribution or morphological defects in the constricting cells. (**A**) Schematic illustrating different scenarios in which defects in contractility can result in invagination defects: ii) decreasing apical myosin in the constricting cells; iii) ablating lateral myosin in the constricting cells (Gracia et al., 2019; John and Rauzi, 2021); iv) optogenetic recruitment of basal myosin in constricting cells (Krueger et al., 2018). (B) Projection of the cross-section view of a representative control and *dlg1* RNAi embryo expressing Sqh-mCherry and E-cadherin-GFP (not shown) imaged with a multiphoton microscope. Basal myosin is downregulated in the *dlg1* RNAi embryo like in the control embryo (yellow arrowheads). T = 0:00 (mm:ss) is the onset of gastrulation. Scale bar: 20 μm. (**C-D**) Confocal images of a representative control and *dlg1* RNAi embryo expressing Sqh-mCherry and E-cadherin-GFP (not shown). Maximum projections of the cross-section view (**C**) and the *en face* view (**D**) are shown. Apical myosin and lateral myosin (red arrows) accumulation are comparable in the control and the *dlg1* RNAi embryos during apical constriction. T = 0:00 (mm:ss) is the onset of gastrulation. Scale bar: 20 μm. (**E**) Confocal images of a representative control and *dlg1* RNAi embryo expressing Sqh-mCherry and E-cadherin-GFP during apical constriciton. The spatial organization of apical myosin in the constricting cells is comparable between the control and *dlg1* RNAi embryos.

### Flanking, non-constricting cells in the dlg1 RNAi embryos exhibit abnormal apical morphology during apical constriction

Since the constricting cells in the *dlg1* RNAi embryos behave relatively normally before the invagination phase, we asked whether the delay in invagination is caused by defects in the non-constricting cells adjacent to the constriction domain. We focused our analysis on the lateral mesodermal cells that connect the constriction domain to the ectodermal cells (Fig. 1A). The lateral mesoderm consists of ∼ 3 rows of cells on each side of the constriction domain. We will refer to these cells as “flanking cells” for the remainder of the text. Before gastrulation, the apical morphology of the flanking cells in the control and *dlg1* RNAi embryos is comparable. Differences between control and *dlg1* RNAi embryos appeared during apical constriction. In control embryos, the apical domain of the flanking cells is moderately stretched along the medial-lateral axis by the neighboring constricting cells, resulting in an extended shape at T_trans_ (Fig. 4A). In *dlg1* RNAi embryos, the apical domain of the flanking cells is stretched to a significantly greater extent than in the control embryos (Fig. 4A). Flanking cells that were extremely stretched in the *dlg1* RNAi embryos gradually developed an abnormal apical morphology over time that became more irregular than their control counterparts (Fig. 4A). To further characterize the overstretched phenotype, we segmented the apical domains of the flanking cells from the surface view of the embryos at T_trans_ and fit each cell apex into an ellipse to determine its area, long axis length, and aspect ratio (long axis/short axis) (Fig. 4B; Methods). The apical domains of the flanking cells in the *dlg1* RNAi embryos are generally more elongated, with an average area of 74.1 ± 27.2 μm^2^ compared to 70.1 ± 23.2 μm^2^ in the control embryos (mean ± s.d., *p* = 0.0003), an average long axis of 14.4 ± 4.0 μm compared to 13.4 ± 3.2 μm in the control embryos (mean ± s.d., *p* = 3 × 10^-10^), and an average aspect ratio of 2.4 ± 0.8 compared to 2.1 ± 0.5 in the control embryos (mean ± s.d., *p* = 2 × 10^-20^) (Fig. 4C; control: 883 cells in 14 embryos; *dlg1* RNAi: 1157 cells in 25 embryos).

**Figure 4.**
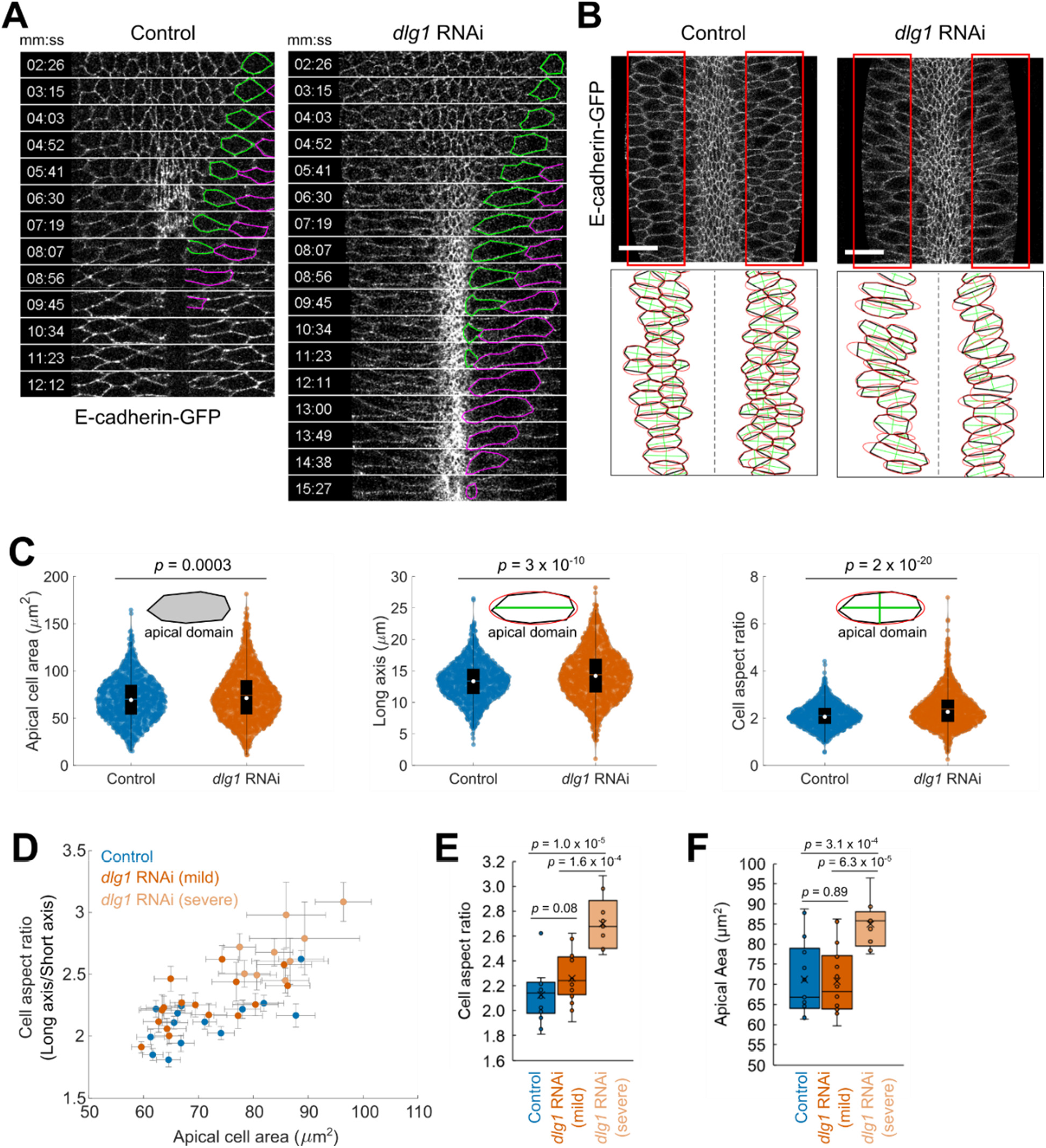
Flanking non-constricting cells in *dlg1* RNAi embryos are over-stretched during apical constriction. **(A)** Time-lapse images of a control embryo and a *dlg1* RNAi embryo during ventral furrow formation. The ventral surface views are shown. In control embryos, the apical domain of the flanking non-constricting cells are moderately stretched before being rapidly internalized. In the *dlg1* RNAi embryos, the flanking non-constricting cells on both sides of the constriction domain are hyper-stretched. T = 0:00 (mm:ss) is the onset of gastrulation. Scale bars: 20 μm. **(B)** Top panels: The surface view of a control embryo and a *dlg1* RNAi embryo expressing E-cadherin-GFP at T_trans_. Bottom panels: The apical cell shape of the flanking non-constricting cells (red boxes in A) at T_trans_ was segmented (black) and fit to an ellipse (red). The long and short axes of the cells (green) were determined based on the fitted ellipses. **(C)** Measurement of geometrical properties of the apical domain of the flanking cells at T_trans_. Left: area; middle: length (long axis); right: aspect ratio. Control: n = 883 cells from 14 embryos. *dlg1* RNAi: n = 1157 cells from 25 embryos. **(D)** The aspect ratio of the flanking non-constricting cells plotted against apical area. Each dot shows the average measurement (n = 52 ± 18 cells per embryo, mean ± std) from a single embryo. Error bars: standard error. Control: n = 14 embryos; mild *dlg1* RNAi: n = 16 embryos; severe *dlg1* RNAi: n = 9 embryos. **(E, F)** Boxplots showing the distribution of the apical cell aspect ratio **(E)** and apical cell area **(F)** in control, *dlg1* RNAi (mild), and *dlg1* RNAi (severe) embryos. Student t-test (unpaired, two-tail) was used for statistical analyses.

Interestingly, we observed a wider range of values for apical area, long axis length, and cell aspect ratio for the flanking cells in the *dlg1* RNAi embryos than in the control embryos (Fig. 4C). We speculated that this was due to embryo-to-embryo variation and differences in knockdown efficiency in the *dlg1* RNAi background. Indeed, we found that the extent of stretching of the flanking cells in the *dlg1* RNAi embryos at T_trans_ varies widely between embryos (Fig. 4D). The variation in the extent of flanking cell stretching that we observed in the *dlg1* RNAi embryos allowed us to test whether the defects in flanking cell morphology are associated with a delay in invagination. Specifically, we asked whether *dlg1* RNAi embryos with mild and severe invagination phenotypes show differences in their flanking cell phenotype. On average, the flanking cells in the mild group (16 out of 25 embryos) and the control group exhibit no significant difference in apical area and apical morphology (Fig. 4D-F). In contrast, the average aspect ratio and apical area of the flanking cells in the severe group is greater than in the control group (aspect ratio: 2.7 ± 0.2 compared to 2.1 ± 0.2 in control embryos, mean ± s.d., *p* = 1.0 × 10^-5^; apical area: 85.0 ± 5.9 μm^2^ compared to 71.2 ± 9.4 μm^2^ in control embryos, mean ± s.d., *p* = 3.1 × 10^-4^; control, n = 14 embryos; severe *dlg1* group: n = 9 embryos) (Fig. 4D-F). The observed correlation between the overstretched phenotype of the flanking cells and the delay in invagination in the *dlg1* RNAi embryos suggests a potential role for the flanking cells in facilitating the transition between apical constriction and invagination.

### Aberrant apical myosin contractions in the flanking cells of dlg1 RNAi embryos contribute to the hyper-stretched phenotype

Next, we sought to determine what causes the flanking cells in the *dlg1* RNAi embryos to become overstretched. It has been recently shown that in addition to the constricting cells, the flanking cells in wildtype embryos also display apical myosin pulses during ventral furrow formation (Denk-Lobnig et al., 2021), as do lateral ectodermal cells during germband elongation (Fernandez-Gonzalez and Zallen, 2011; Rauzi et al., 2010; Sawyer et al., 2011). In the constricting cells, the apical actomyosin network undergoes ratcheted contractions that result in a stepwise reduction of cell apical area. In contrast, the flanking cells accumulate lower levels of apical myosin, which result in unratcheted myosin pulses that do not lead to a net apical area reduction. We hypothesized that the unratcheted myosin pulses in the flanking cells allow the cells to temporarily resist stretching induced by pulling forces from the constriction domain. We further hypothesized that this mechanism is no longer functional in the *dlg1* RNAi embryos, thereby leading to overstretching of the flanking cells.

To test these hypotheses, we first examined the relationship between myosin activity and apical cell dynamics in control flanking cells. Consistent with a previous report (Denk-Lobnig et al., 2021), we detected apical myosin coalescence in the flanking cells as apical constriction progressed (Fig. 5A; Movie 4). Compared to the constricting cells, accumulation of apical myosin in the flanking cells occurred later, and apical myosin intensity was much lower (Fig. S5A; cyan arrows). Typically, we were able to observe several rounds of myosin coalescence in an individual flanking cell before it disappeared from the surface of the embryo (Fig. S5B; red asterisks). In order to quantify apical myosin activity in the flanking cells and analyze its impact on apical cell shape dynamics, we segmented the apical domain of individual flanking cells and displayed apical myosin intensity within individual cells over time as heatmaps. We focused our analysis between 4 minutes before T_trans_ and 2 minutes after T_trans_ (approximately between 4 – 10 minutes after the onset of apical constriction), when myosin coalescence was most frequently observed in the flanking cells. Consistent with the observed cycles of myosin coalescence, the heatmaps revealed the repeated rise and fall of myosin intensity in the flanking cells, with a general trend of intensity increase over time (Fig. 5B). To analyze the impact of myosin coalescence on apical cell shape changes, we identified myosin pulses by manually determining pulse peak time, which we defined as the time when myosin coalesced into a high-intensity punctum at the apical domain (Fig. 5C-D; Time 00:00). We then asked how apical cell area changes during a 96-second interval centered around the pulse peak time. The 96-second interval was selected because it usually covers the rising and falling phases of myosin intensity during a single coalescence event, meanwhile minimizing overlap between consecutive pulses. These 96-second intervals are referred to as “myosin pulses” hereafter (examples shown in Fig. 5B; red boxes). We found that myosin pulses in control flanking cells are often, albeit not always, accompanied by a reduction in cell area (Fig. 5C, E; Fig. S6A). We reasoned that the weak correlation between myosin pulses and cell area reduction is likely due to the impact of pulling forces from the constriction domain, which was not accounted for in our analysis. To test whether the observed myosin pulses have any impact on apical cell area in the flanking cells, we compared apical cell area change during myosin pulses and during 96-second intervals between two successive myosin pulses (“off-pulses”, Methods). During a pulse, apical myosin generally reaches its highest intensity at the pulse peak time (Fig. 5G). As expected, apical myosin intensity at the pulse peak time was significantly higher during pulses than during off-pulses (Fig. 5H). Interestingly, we found that the average apical cell area change differed between pulses and off-pulses. During pulses, the average apical area barely changed, although there was a slight reduction in apical area at the pulse peak. In contrast, apical cell area increased significantly during off-pulses (Fig. 5I-J; apical area change was -0.4 ± 11.8 μm^2^ for pulses and 7.1 ± 12.1 μm^2^ for off-pulses; n = 56 pulses and 41 off-pulses, *p* = 0.0018). The change in length of the apical domain along the medial-apical axis (“apical length”) displayed a similar trend as the change in apical cell area (Fig. 5K-L; apical length change was 1.1 ± 2.3 μm for pulses and 2.0 ± 2.3 μm for off-pulses; n = 56 pulses and 41 off-pulses, *p* = 0.036). Note that these differences are not due to initial differences in cell size and morphology at the beginning of pulses, as a similar trend was observed when we normalized apical area and apical length by their initial size (Fig. S6B). Together, this data supports a model in which myosin pulses in the flanking cells function to restrain stretching of the apical domain, presumably by counteracting pulling forces from the constriction domain.

**Figure 5.**
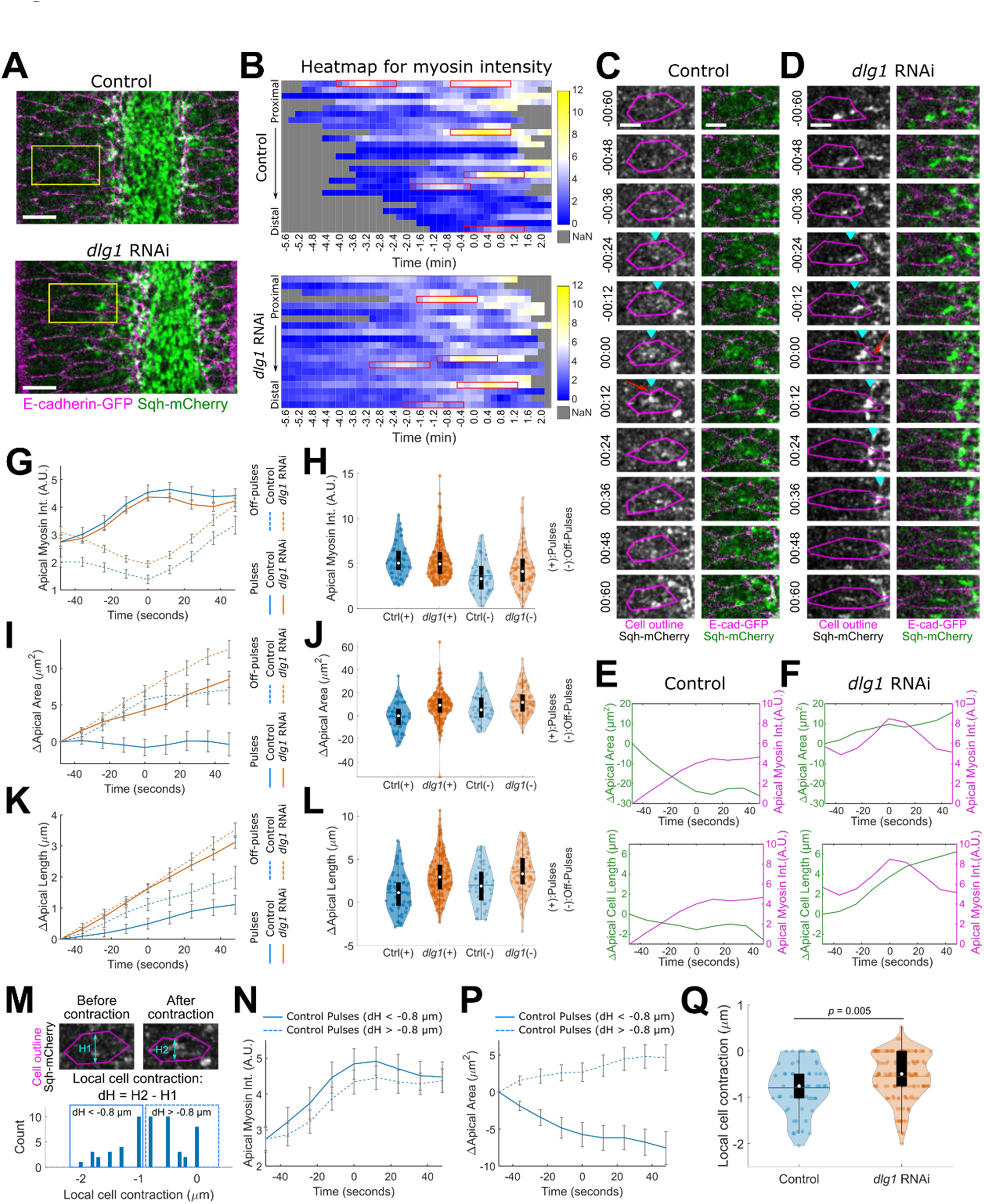
Knockdown of Dlg1 affects apical myosin contractions in the flanking cells. **(A)** Ventral surface view of a control embryo and a *dlg1* RNAi embryo expressing E-cadherin-GFP and Sqh-mCherry at the end of the apical constriction phase. The flanking cells highlighted by yellow boxes are presented in panel C and D. Scale bars: 10 μm. **(B)** Heatmap showing apical myosin intensities over time in individual flanking cells. Each row corresponds to one cell. The cells are arranged based on their proximity to the ventral midline. T = 0:00 (mm:ss) corresponds to T_trans_. Gray indicates time points for which data is unavailable. **(C, D)** Examples of apical myosin coalescence observed in the flanking cells in control and *dlg1* RNAi embryos. T = 0:00 (mm:ss) corresponds to the time when myosin intensity is the highest for an individual pulse (the pulse peak). Both cells are on the left side of the ventral midline. Myosin coalescence is indicated by cyan triangles. Local cell contractions following myosin pulses are indicated by red arrows. Scale bars: 10 μm. **(E, F)** Apical myosin intensity and changes in apical cell area and apical cell length over time in cells shown in C and D, respectively. Time zero is the pulse peak. **(G, I, K)** Average trend of apical myosin intensity, apical area change, and apical length change over time for pulses and off-pulses in the flanking cells. 56 pulses and 41 off-pulses from 4 control embryos and 143 pulses and 100 off-pulses from 8 *dlg1* RNAi embryos were analyzed. **(H)** Distribution of apical myosin intensity at the pulse peak. **(J, L)** Distribution of apical area change and apical length change over a pulse or over an off-pulse in the flanking cells. **(M)** Top: Schematic showing measurement of local cell contraction (dH) during a pulse in the flanking cells. Bottom: Distribution of dH for pulses in the control embryos. **(N, P)** Average trend of apical myosin intensity and apical area change over time for control pulses associated with high (dH < -0.8 μm) or low (dH > -0.8 μm) levels of local cell contraction. **(Q)** Distribution of dH for pulses in control and *dlg1* RNAi embryos.

Next, we performed a similar analysis in the *dlg1* RNAi embryos. Similar to the control flanking cells, the mutant flanking cells also displayed cycles of myosin pulses that had a similar duration and frequency (Fig. 5B; Fig. S6C). Furthermore, we found that the average apical myosin intensity during pulses is comparable between the two genotypes (Fig. 5G-H). Despite these similarities, myosin pulses in the flanking cells of *dlg1* RNAi embryos are rarely associated with either reduction or stabilization of apical cell area or apical length (Fig. 5D, F; Fig. S6A). The average increase in apical cell area and apical length during pulses in the mutant flanking cells is significantly higher than in the control flanking cells and appears to be more similar to non-pulses (Fig. 5I-L; apical area change was 8.5 ± 13.2 μm^2^ for *dlg1* pulses and 12.6 ± 12.0 μm^2^ for *dlg1* off-pulses; n = 143 pulses and 100 off-pulses, *p* = 5.6 × 10^-6^ for comparison between control and *dlg1* pulses; apical length change was 3.1 ± 2.4 μm for *dlg1* pulses and 3.5 ± 2.3 μm for *dlg1* off-pulses; 1.4 × 10^-7^ for comparison between control and *dlg1* pulses). These observations suggest that myosin pulses in the mutant flanking cells are much less effective at restraining apical cell stretching than those in control flanking cells.

### Coupling between apical myosin coalescence and local cell deformation is weakened in the flanking cells of dlg1 RNAi embryos

What causes the difference in the effect of myosin pulses on local cell shape changes between control and *dlg1* mutant flanking cells? A closer examination of myosin pulses in control flanking cells suggests that there are various degrees of coupling between myosin coalescence and local cell shape deformation. When coupling is strong, myosin coalescence correlates with pulling and bending of the cell-cell boundary towards the center of the apical domain (Fig. 5M). When there is weak or no coupling, no obvious local deformation is observed during myosin coalescence. To further analyze these behaviors, we measured the local width change of the apical domain during the rising phase of the pulse at the location where apical myosin coalescence occurs (“dH”; Fig. 5M). We used dH as the “coupling index” to indicate the degree of coupling. To validate this approach, we compared apical cell area change during strongly coupled pulses and weakly coupled pulses in control embryos (dH < -0.8 μm and dH > - 0.8 μm, respectively; Fig. 5M). We found that during strongly coupled pulses, the apical area of the flanking cells undergoes a net reduction. In contrast, apical area increased during weakly coupled pulses (Fig. 5N, P). The result is consistent with the expectation that more strongly coupled pulses result in more prominent apical area reduction. Next, we measured the coupling index, dH, for the pulses in the *dlg1* RNAi mutants. The average dH was significantly less (i.e., closer to zero) for pulses in the *dlg1* RNAi mutants than in the control embryos, indicating that there is less change in cell width per pulse (Fig. 5Q; dH was -0.79 ± 0.55 μm for control pulses and -0.55 ± 0.5 μm for *dlg1* pulses; *p* = 0.005). These results suggest that coupling between myosin coalescence and local cell deformation is reduced in the mutant flanking cells. Together, our results suggest that reduced coupling between contractile myosin and cell-cell boundaries in the mutant flanking cells impairs their ability to resist cell stretching.

The cause of weaker coupling between myosin coalescence and local cell deformation upon Dlg1 depletion is unknown. Interestingly, we noticed that the adherens junction marker E-cadherin-GFP shows reduced intensity at the subapical region in the mutant flanking cells (Fig. S7). In the constricting mesodermal cells, the apical myosin network exerts contractile forces at cell-cell boundaries by anchoring at the apical adherens junctions (Martin et al., 2010). The observed defect in apical adherens junctions in the mutant flanking cells led us to hypothesize that the “unclutched” myosin pulses are the result of impaired actomyosin fiber attachments at the cell-cell junctions. Consistent with this hypothesis, we frequently observed rapid flow of myosin to the leading edge (the edge closer to the midline) of the flanking cells in the *dlg1* RNAi embryos (Fig. 5D; cyan triangles). This phenomenon can be explained by reduced anchoring of actomyosin at cell-cell boundaries (Fig. 5D).

Based on these observations, we propose that the overstretched phenotype of the flanking cells in the *dlg1* RNAi embryos is at least partially caused by defects in coupling between apical myosin and adherens junctions. In control flanking cells, the contractile myosin structures are coupled to the cell-cell boundaries. The resulting clutched myosin contractions are able to counteract the pulling forces from the constriction domain, and thereby restrain stretching of the flanking cells. In the mutant flanking cells, the fraction of “unclutched” myosin pulses increases, which impairs the ability of the flanking cells to resist pulling from the constriction domain and results in hyper-stretching.

### Knockdown of Dlg1 affects how cells respond to ectopic pulling forces prior to gastrulation

We noticed that the extent of apical area increase in the flanking cells during off-pulses is larger in the *dlg1* RNAi embryos than in the control embryos (Fig. 5I, K), suggesting that in addition to unclutched myosin pulses, other factors may also contribute to the overstretched phenotype. For example, knockdown of Dlg1 may alter the “passive” mechanical properties of cells, making them more deformable. To test this possibility, we used a magnetic tweezers-based approach to probe tissue mechanical properties in live embryos shortly before gastrulation. In these experiments, we injected magnetic beads into the ventral-lateral region of embryos during early cellularization. These beads became enclosed in the newly formed cells, which allowed us to exert forces on the tissue using an electromagnet and record the response simultaneously (Fig. S8A-B; Methods). Since the degree of tissue deformation during the pulling phase is subject to variation from the number of beads injected into the embryo, we focused our analysis on a group of control and *dlg1* RNAi embryos that had a comparable number of beads and a similar degree of tissue deformation (Fig. S8F). In control embryos, the tissue undergoes viscoelastic recoil after release from pulling (Fig. S8C-E). In *dlg1* RNAi embryos, the tissue also displayed recoil after pulling, but the degree of recoil was reduced compared to that observed in the control (Fig. S8F-H). This observation suggests that the cells in the *dlg1* RNAi embryos are slightly more viscous and less elastic than those in the control embryos. Due to the limited accuracy and precision of bead injection, we were not able to specifically measure the response of the flanking cells. Nevertheless, the difference between the control and *dlg1* RNAi cells in their response to ectopic pulling forces raises the interesting possibility that knockdown of Dlg1 may result in changes in tissue mechanical properties that contribute to the overstretched phenotype of the flanking cells.

### Hyper-stretching of the flanking cells in the dlg1 RNAi embryos slows down the ventral movement of the ectodermal tissue during ventral furrow formation

During ventral furrow formation, the neighboring ectodermal tissue moves towards the ventral midline as the mesoderm internalizes. Since the constriction domain and the ectodermal tissue are connected by the flanking cells, we predicted that hyper-stretching of the flanking cells in the *dlg1* RNAi embryos would delay ventral movement of the ectodermal tissue (Fig. 6A). To test this prediction, we generated surface projections of the embryos from the multiphoton movies described in Figure 2. These apical surface views covered a wide region of the embryo along the medial-lateral axis, which allowed us to examine the displacement of the constricting mesodermal cells, the flanking cells, and the neighboring ectodermal cells at the same time.

**Figure 6.**
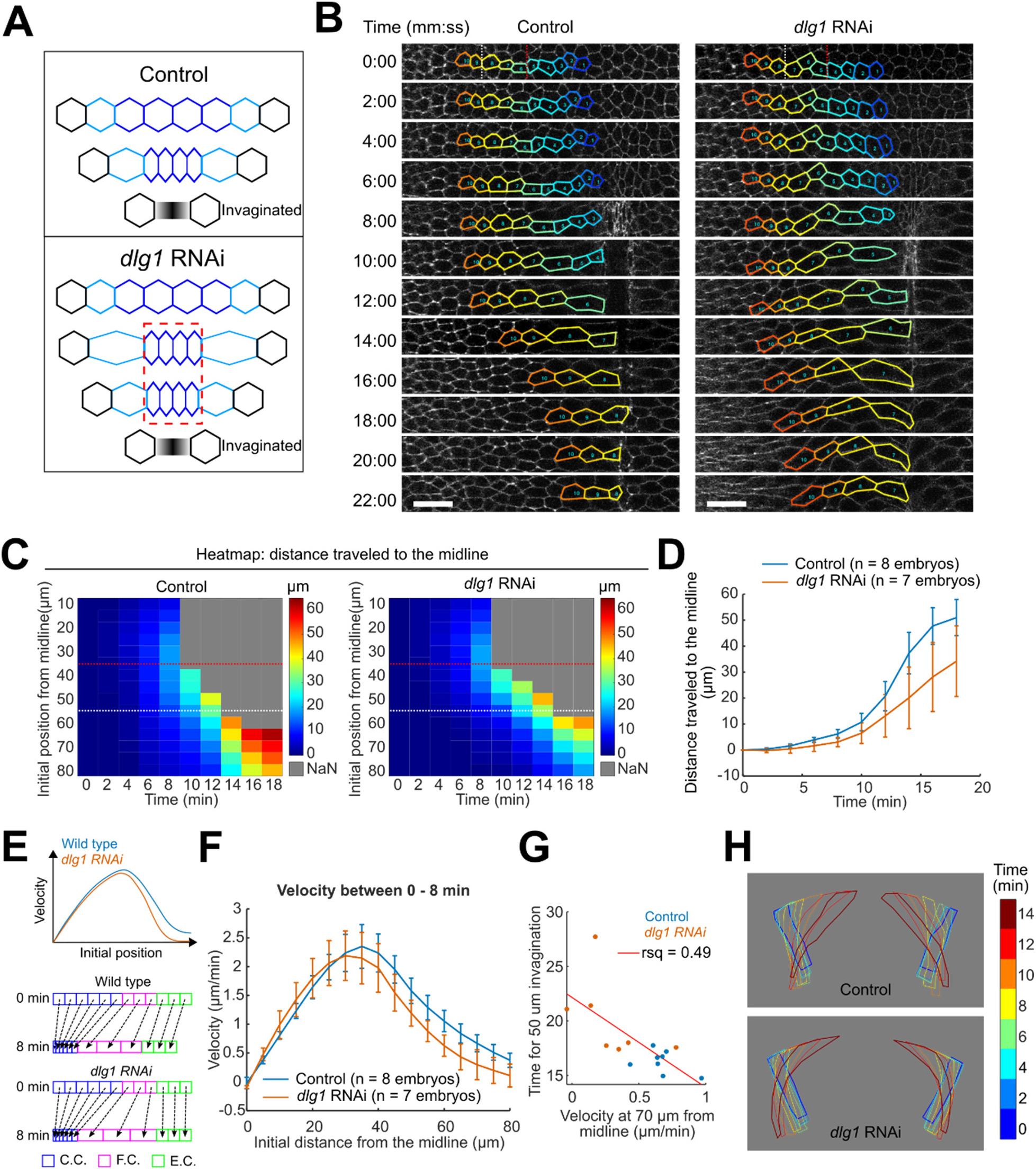
Ventrally directed movement of the lateral ectodermal cells is delayed in *dlg1* RNAi embryos. **(A)** Schematic illustrating the shape change of the apical domain of the constricting cells (dark blue) and the flanking cells (light blue) during apical constriction. The dashed red box highlights the extended transition phase in the *dlg1* RNAi embryos that is associated with the hyper-stretched phenotype of the flanking cells. Hyper-stretching of the flanking cells is expected to delay the ventral movement of the ectodermal cells (black). **(B)** Movie stills showing the movement and shape change of cells (outlined) initially located at different locations along the medial-lateral axis on the left side of the ventral midline. The surface view of the embryos is shown. Each color represents one cell that has been tracked over time during ventral furrow formation. T = 0:00 (mm:ss) is the onset of gastrulation (same for all other panels). At the onset of gastrulation, the constricting cells, the flanking cells, and the neighboring ectodermal cells are distributed approximately 0 – 35 μm, 35 – 55 μm, and more than 55 μm from the ventral midline, respectively. The red dotted line indicates the boundary between the constricting and flanking cells, while the white dotted line indicates the boundary between the flanking cells and the ectodermal cells. Scale bars: 20 μm. **(C)** Heatmaps showing the distance that cells traveled towards the ventral midline over time that are initially located at different medial-lateral positions. Two example embryos are shown. The red dotted line indicates the boundary between the constricting and flanking cells, while the white dotted line indicates the boundary between the flanking cells and the ectodermal cells. **(D)** Distance traveled towards the midline for ectodermal cells initially located 70 μm away from the midline. **(E)** Schematic showing cell velocity during apical constriction as a function of the initial medial-lateral position of cells located at different distances from the ventral midline. C.C.: constricting cells; F.C.: flanking cells; E.C., ectodermal cells. Hyper-stretching of the flanking cells is expected to reduce the velocity of ectodermal cell movement. **(F)** Measurement of cell velocity during apical constriction as a function of the initial medial-lateral position of cells located at different distances from the ventral midline. **(G)** A reduced rate of ectodermal cell movement during apical constriction correlates with invagination delays. **(H)** The cross-section view of a 2D outline of two flanking cells over time in a representative control and *dlg1* RNAi embryo.

For each embryo, we segmented the apical cell outlines of a single row of cells along the medial-lateral axis and tracked them over time (Fig. 6B). In the *dlg1* RNAi embryos, hyper-stretching of the flanking cells mainly occurs in the apical domain, which results in increased bending of the flanking cells towards the ventral midline (Fig. 6H). To examine how overstretching of the flanking cells affects coordination of tissue movement during ventral furrow formation, we measured cell displacement over time (Fig. 6C). Movement of the ectodermal cells towards the ventral midline was substantially slower in the *dlg1* RNAi embryos, as indicated by the shorter distance traveled towards the ventral midline (Fig. 6C-D). In theory, both overstretching of the flanking cells and delayed mesoderm invagination could result in a reduced rate of ectodermal movement. We therefore wondered whether the defects in ectodermal cell movement in the *dlg1* RNAi embryos could be detected prior to T_trans_, before invagination of the mesoderm occurs. As illustrated in Figure 6E, prior to T_trans_, we predicted that hyper-stretching of the flanking cells in the *dlg1* RNAi embryos would mostly affect the rate of ectodermal cell movement, and to a lesser extent, flanking cell movement. Our measured velocities of cell movement during the first 8 minutes of ventral furrow formation closely matched the predicted outcome (Fig. 6F). Together, these analyses indicate that overstretching of the flanking cells results in a delay in ectodermal movement that can be detected before the onset of rapid invagination. Notably, there is a moderate correlation between the rate of ectodermal cell movement during apical constriction and the delay in invagination (Fig. 6G). This result led us to hypothesize that proper coupling between the constricting cells and the ectodermal tissue is important for a robust transition between apical constriction and invagination during ventral furrow invagination.

### Laser ablation-mediated disruption of the flanking cells delays the transition between apical constriction and invagination

The phenotypic analysis of *dlg1* RNAi embryos prompted us to ask whether disrupting the flanking cells is sufficient to cause invagination defects. To test this, we used a focused beam of a near-infrared laser to disrupt the flanking cells in wildtype embryos during apical constriction. To avoid any adverse effects from wound healing responses, we carefully tuned the laser power such that no visible damage on the plasma membrane was observed after laser ablation. Prompt recovery of the plasma membrane signal of E-cadherin-GFP after laser ablation suggests that the plasma membrane was not damaged (data not shown). However, we cannot confidently conclude that the plasma membrane was not perturbed at all, as there could have been damage that was not detectable under the imaging conditions we used. We confirmed the effectiveness of this approach by treating tissues that were under tension. After laser ablation, we observed an immediate retraction of the surrounding tissues from the ablated site, an expected tissue response when the mechanical integrity of a laser-treated region is disrupted (data not shown).

Next, we used this approach to disrupt the flanking cells, which we expected would weaken mechanical coupling between the constricting cells and the neighboring ectodermal tissue. We scanned a focused laser beam across a ∼170 μm × 10 μm region at the apical surface spanning approximately 2 columns of flanking cells on both sides of the constriction domain (Fig. 7A-B). We performed laser ablation approximately 2-3 minutes before T_trans_, when the flanking cells are stretched and are expected to be under tension. After laser ablation, the ablated region immediately expanded and the non-constricting tissue lateral to the treated region underwent immediate retraction (Fig. 7C, cyan arrows; Movie 5). This observation confirms that the flanking cells are under tension during apical constriction and further indicates that the ectodermal cells are pulled on by the flanking cells. We did not observe any obvious impacts on the constriction domain after laser ablation – no relaxation happened, and the cells continued to constrict apically (Fig. 7C, magenta arrows; Movie 5). Interestingly, we noticed that approximately one minute after laser ablation, the treated region that underwent expansion started to shrink, and coupling between the constriction domain and the ectoderm appeared to be restored (data not shown). The reason why the ablated region shrank is unclear. In order to prevent recoupling of the constriction domain and the ectoderm, we repeated laser ablation every 68 seconds to achieve prolonged disruption of the flanking cells. Under these conditions, we examined the consequence of flanking cell disruption during apical constriction on ventral furrow formation. In “non-ablated” embryos, the neighboring ectodermal cells moved persistently towards the ventral midline and the ventral furrow invaginated normally (Fig. 7C-F). In the “ablated” embryos, movement of the neighboring ectodermal cells towards the ventral midline was interrupted, even though the laser treatment did not noticeably affect the constriction domain (Fig. 7C; Movie 5). Most interestingly, this interruption was associated with a delay in the transition between apical constriction and invagination (Fig. 7D-F; Movie 5). In all cases, ventral movement of the ectodermal cells and invagination of the ventral furrow eventually resumed after a certain delay (Fig. 7D-E). It is unclear what lead to the recovery of invagination, but we speculate that it was facilitated by active movement of the ectodermal tissue towards the ventral midline, which occurs at later stages of ventral furrow formation (see Discussion). Nevertheless, our results demonstrate that decoupling of the constricting mesodermal cells and the nearby ectodermal tissue results in a delay in invagination, which is consistent with the phenotype observed in the *dlg1* RNAi embryos. This result supports the hypothesis that coupling between the constriction domain and the ectodermal tissue is important for an effective transition between apical constriction and invagination.

**Figure 7.**
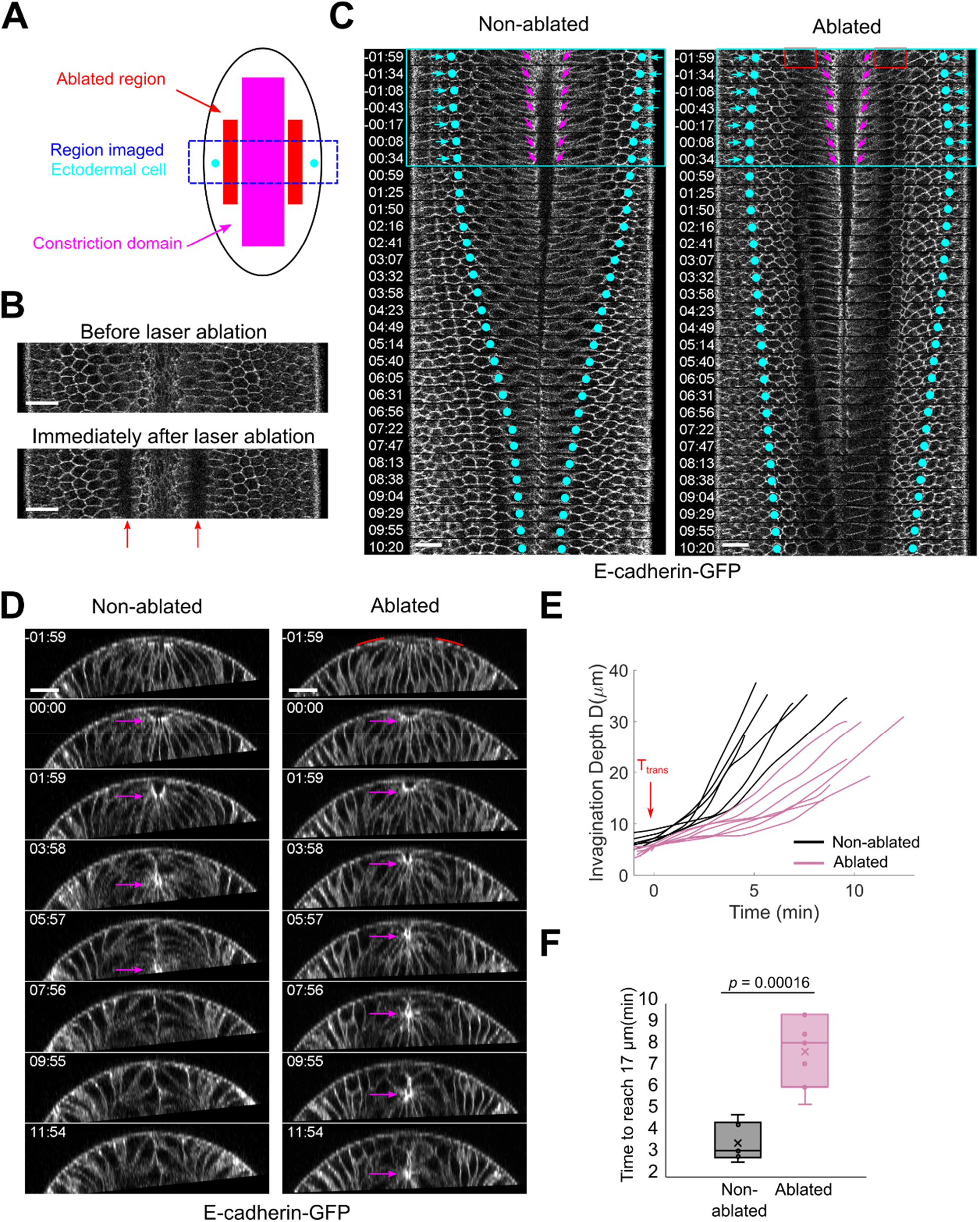
Laser ablation of flanking non-constricting cells during apical constriction in wildtype embryos delays the transition between apical constriction and invagination. (**A**) Schematic illustrating the laser ablation experimental design in a *Drosophila* embryo. A row of flanking cells was ablated by a laser (red box). A portion of the embryo (blue box) was imaged before and after ablation. The magenta box indicates the position of the constriction domain, and the cyan dots demarcate the position of two ectodermal cells on opposite sides of the ventral midline. (**B**) Ventral surface view of an embryo before (top) and after (bottom) laser ablation. Red arrows indicate the ablated regions. Scale bars: 20 μm. (**C-D**) Representative *en face* (**C**) and cross-section (**D**) images of stage matched wildtype embryos undergoing ventral formation with or without laser ablation. T = 00:00 (mm:ss) corresponds to the transition from apical constriction to invagination in control embryos. Cyan dots in C indicate the position of two ectodermal cells on opposing sides of the ventral midline. Note that the ectodermal cells temporarily underwent retraction following laser ablation (cyan box and arrows), suggesting that they were pulled on by the flanking cells before laser ablation. The constriction domain appears to be relatively unaffected by laser ablation, as illustrated by the magenta arrows in C. Magenta arrows in D indicate the position of the apex of the constricting cells. Scale bars: 20 μm. (**E**) Quantification of ventral furrow invagination depth, D, over time in non-ablated (n = 6) and ablated (n = 7) embryos. (**F**) Boxplots showing the time it takes for the ventral furrow to reach 17 μm in non-ablated and ablated embryos. Student t-test (unpaired, two-tail) was used for statistical analysis.

### Acute disassembly of the cortical actin cytoskeleton in the flanking cells delays the transition between apical constriction and invagination

To further test whether the mechanical integrity of the flanking cells is important for facilitating tissue invagination, we used a previously developed CRY2-CIBN optogenetic tool to downregulate actin in the flanking cells during apical constriction (Guglielmi et al., 2015). Cryptochrome 2 (CRY2) is a blue light absorbing photosensor that binds to the N-terminal domain (CIBN) of the transcription factor, CIB1, in its photoexcited state (Liu et al., 2008b). Since previous studies have shown that optogenetic downregulation of actin in the constricting cells impairs myosin contractility during ventral furrow formation (Guglielmi et al., 2015), we reasoned that actomyosin contractions and resistance in the flanking cells would also be disrupted after targeted stimulation. In this optogenetic system, CIBN is anchored to the plasma membrane (CIBN-pm). Upon light stimulation, CIBN recruits cytoplasmic CRY2-OCRL (tagged with mCherry) to the plasma membrane (Fig. 8A-B). OCRL is the catalytic domain of the inositol polyphosphate 5-phosphatase (Zhang et al., 1995). Once at the plasma membrane, OCRL converts PI(4,5)P_2_ into PI(4)P, which results in disassembly of cortical actin. Previous studies have shown that precise stimulation (confined within a focal volume) can be achieved with 920 nm multiphoton laser illumination (Guglielmi et al., 2015).

**Figure 8.**
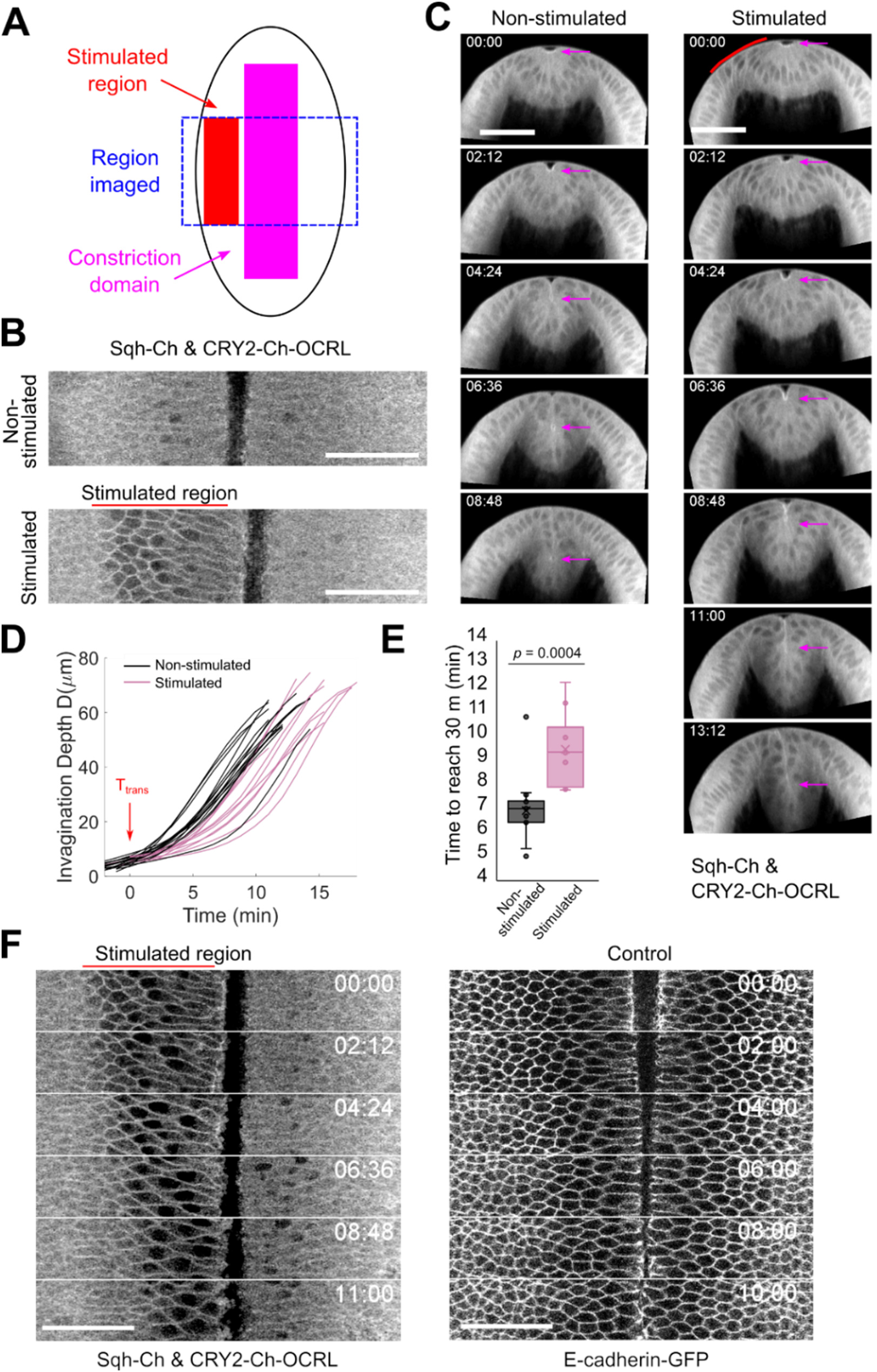
Downregulation of cortical F-actin in the lateral non-constricting cells delays the transition between apical constriction and invagination. (**A**) Schematic illustrating the optogenetic experimental design in a *Drosophila* embryo. Embryos are stimulated during apical constriction in a confined region spanning 4 – 5 rows of non-constricting cells on one side of the constriction domain (red box). A portion of the embryo (blue box) was imaged before and after stimulation. The magenta box indicates the position of the constriction domain. (**B-C**) Representative surface view (**B**) and cross-section (**C**) images of stage matched embryos expressing CIBN-pm-GFP and CRY2-mCherry-OCRL undergoing ventral furrow formation in the presence or absence of stimulation. The cytoplasmic localization of CRY2-mCherry-OCRL in the absence of stimulation and the plasma membrane recruitment of CRY2-mCherry-OCRL upon stimulation can be observed from the surface view in B. T = 0:00 (mm:ss) corresponds to the transition from apical constriction to invagination in control embryos. The red lines highlight the stimulated region. Scale bars: 50 μm. (**D**) Quantification of ventral furrow invagination depth, D, over time in the absence of stimulation (n = 15 embryos) and in the presence of stimulation (n = 10 embryos). (**E**) Boxplots showing the time it takes for the ventral furrow to reach 30 μm in non-stimulated and stimulated embryos. Student t-test (unpaired, two-tail) was used for statistical analysis. (**F**) Movie stills showing the surface view of a representative stimulated embryo expressing CIBN-pm-GFP and CRY2-mCherry-OCRL (left) and the surface view of a representative control embryo expressing E-cadherin-GFP but not the optogenetic constructs (right). Optogenetic depletion of F-actin results in abnormal cell shape in the stimulated region of embryos (left), which is not observed in control embryos (right).

In order to test the effectiveness of the system, we performed multiphoton stimulation on a single apical plane encompassing half of an embryo expressing Utr-Venus, Sqh-mCherry, and the optogenetic OCRL constructs. After 2 minutes of stimulation, we examined the response of cortical actin over time. Significant actin downregulation was observed immediately after multiphoton stimulation in the stimulated region, but not in the unstimulated region, confirming that multiphoton stimulation is spatially precise (Fig. S9). No obvious actin signal recovery is detected 8-10 minutes after stimulation in the stimulated region (data not shown), which is consistent with previous reports (Guglielmi et al., 2015).

After validating the optogenetic system, we first examined ventral furrow formation in embryos expressing the optogenetic OCRL constructs in the absence of light stimulation. Images were acquired using the 1040 nm laser, which does not stimulate CRY2-OCRL (Guglielmi et al., 2015). In the absence of light stimulation, ventral furrow formation proceeds relatively normally in most embryos. A small percentage of embryos (3 out of 18 embryos examined) were morphologically abnormal during apical constriction and failed to invaginate. Abnormal embryos that showed defects in apical constriction were not included in our analysis. Next, we tested whether stimulation of the flanking cells during apical constriction would affect furrow invagination. Since it took significantly longer to disrupt the flanking cells with optogenetics than with laser ablation, we found it technically more challenging to reliably target the flanking cells on both sides of the constriction domain, like in the laser ablation experiment. We therefore performed stimulation in the flanking cells on one side of the constriction domain. For stimulation, a single apical plane within the selected region of interest was scanned for about 2 minutes using the 920 nm laser. Images before and after stimulation were acquired using the 1040 nm laser. First, we tested stimulation of a 170 μm × ∼ 10 μm rectangular region that spanned one column of flanking cells. Unexpectedly, the treatment resulted in minor effects on invagination, perhaps due to relatively mild disruption of the flanking cells (data not shown). We then increased the width of the stimulated area to ∼ 30 μm, which spanned four to five columns of non-constricting cells next to the constriction domain. The stimulated region mostly covered the flanking cells but also included about 1 – 2 columns of adjacent ectodermal cells (Fig. 8A-B). We found that stimulating this region during apical constriction consistently results in a delay in furrow invagination (Fig. 8C-E; Movie 6). The apical domain of the stimulated cells became more elongated and adopted a more irregular shape than cells in wildtype embryos (Fig. 8F). This behavior is reminiscent of the flanking cells in the *dlg1* RNAi embryos. Taken together, the laser ablation and optogenetic experiments in the wildtype embryos provide additional evidence supporting the notion that maintaining the mechanical integrity of the flanking cells is important for an efficient transition between apical constriction and invagination.

## Discussion

Apical constriction is an important mechanism that promotes folding of flat epithelia in a variety of tissue morphogenetic processes. However, how apical constriction results in tissue folding is not fully understood. Using a combination of genetic, live-imaging, and biophysical approaches, we tried to identify factors other than apical constriction that also influence tissue folding. We present evidence that during *Drosophila* ventral furrow formation, the integrity of the flanking cells, the non-constricting cells adjacent to the constriction domain, is important for a robust transition between apical constriction and invagination. In our search for genes that regulate the invagination phase of ventral furrow formation, but not the apical constriction phase, we identified Dlg1. Dlg1 is a basolateral polarity determinant in epithelial cells. In *dlg1* RNAi embryos, apical myosin activation and apical constriction occur normally, but invagination is defective. Strikingly, the mutant embryos display a prolonged delay in the transition between apical constriction and invagination. Prior to this transition, the morphology of individual constricting cells and their spatial arrangement in the intermediate furrow is largely normal, with the exception that the apical-basal thickness of the intermediate furrow is slightly reduced. Two additional processes that have been recently shown to be important for effective invagination, including the accumulation of myosin along the lateral cortex of the constricting cells and the dissociation of myosin from the basal cortex of the constricting cells (Gracia et al., 2019; John and Rauzi, 2021; Krueger et al., 2018; Polyakov et al., 2014), occur normally in the *dlg1* RNAi embryos. In contrast to the minor defects we observed in the constriction domain, we found prominent morphological defects in the flanking cells of the *dlg1* RNAi embryos. The apical domain of the flanking cells becomes substantially overstretched as the ventral mesodermal cells undergo apical constriction. This overstretching is associated with increased bending of the flanking cells towards the ventral midline. The severity of the overstretched phenotype correlates with the extent of delay in invagination, raising the possibility that appropriate cell shape changes in the flanking cells are important for a smooth transition between apical constriction and invagination. Using two separate approaches, we directly tested the role of the flanking cells in wildtype embryos during ventral furrow formation. In the first approach, we used a focused two-photon laser beam to cut, and thus, impair the mechanical integrity of the flanking cells during apical constriction. In the second approach, we utilized a previously published optogenetic tool to downregulate F-actin in the flanking cells and ∼2 rows of adjacent ectodermal cells (Guglielmi et al., 2015). Importantly, in both treatments, the constricting cells were not perturbed. Both manipulations resulted in a delay in invagination, much like in the *dlg1* RNAi embryos. While we could not directly test the causal relationship between the flanking cell phenotype and the delay in invagination in the *dlg1* RNAi embryos, the observed defects in invagination upon disruption of the flanking cells in otherwise wildtype embryos provides direct evidence that the properties of the flanking cells can influence the tissue folding process, even when apical constriction is normal.

The way by which the flanking cells contribute to invagination remains unclear. Several studies have shown that manipulation of the surrounding non-constricting tissue can impair ventral furrow formation. One study has shown that preventing the movement of the lateral ectoderm, by anchoring ectodermal cell apices to the vitelline membrane, blocks ventral furrow invagination (Rauzi et al., 2015). Another study has shown that upregulation of apical myosin contractility in the lateral ectodermal tissues can reverse the furrow invagination process (Perez-Mockus et al., 2017). In both cases, the invagination defects could be attributed to increased resistance to tissue motion associated with mesoderm invagination. The defects of the flanking cells induced/observed in our study are unlikely to cause an increase in resistance to tissue motion during ventral furrow formation, since we did not observe any obvious tissue anchoring or ectodermal contraction. Instead, we found that defects in the flanking cells result in a delay in ventrally directed movement of the ectodermal cells during apical constriction and invagination. In our laser ablation experiment, we observed immediate retraction of the neighboring ectodermal cells away from the ventral midline upon flanking cell cutting, suggesting that the flanking cells are important for coupling the movements of the constricting cells and the surrounding ectodermal tissue. When the flanking cells are impaired, they are less capable of resisting pulling from the constriction domain and become overstretched. This in turn weakens coupling between apical constriction in the mesoderm and ventrally directed movement of the ectoderm (Fig. 9). Since a delay in the transition between apical constriction and invagination was observed in all three scenarios in which the flanking cells were impaired (i.e., knockdown of Dlg1, laser ablation of the flanking cells, and downregulation of F-actin in the flanking cells), we propose that tissue-level coupling between the mesoderm and the ectoderm is important for promoting mesoderm invagination. How this coordination promotes tissue folding remains to be elucidated. Interestingly, it has been previously shown that ventral movement of the ectoderm is not merely a consequence of mesoderm invagination. It can still occur, albeit with a reduced speed, in mutants where apical constriction is completely abolished (Rauzi et al., 2015). Therefore, it is attractive to propose that ectodermal movement plays an active role in promoting the transition between apical constriction and invagination, perhaps by providing pushing forces on both sides of the ventral furrow. In such a scenario, appropriate coupling between the mesoderm and the ectoderm during apical constriction may facilitate mesoderm invagination by enabling transmission of forces from the ectoderm to the mesoderm. It remains unclear whether these forces exist, and if so, whether they originate from the lateral ectoderm or the dorsal ectoderm. Future experiments that measure and/or alter mechanical forces in the ectoderm during ventral furrow formation will be necessary to test this hypothesis.

**Figure 9.**
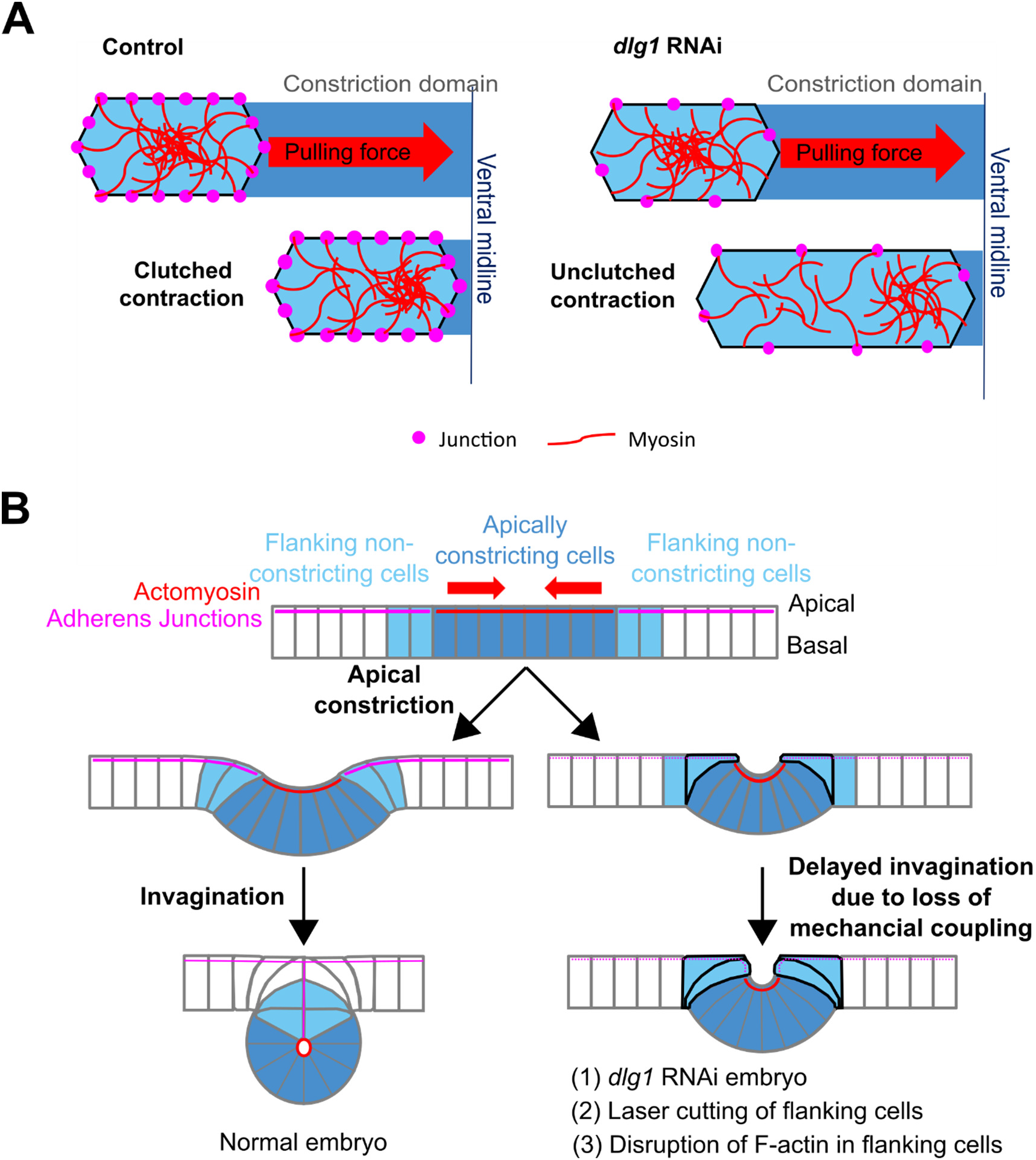
A working model for cellular mechanics and tissue coordination during epithelial folding. **(A)** In control embryos, effective force transmission between the actomyosin network and the adherens junctions allows the flanking cells to undergo clutched contractions. Clutched contractions enable the flanking cells to resist cell shape changes when they are pulled on by the constricting cells. In the *dlg1* RNAi embryos, the flanking cells are also able to accumulate apical myosin, but myosin contractions become “unclutched” and are ineffective in exerting forces on the cell boundary. As a result, the mutant flanking cells are unable to resist pulling from the constricting cells and become hyper-stretched. We hypothesize that the weakened adherens junctions we observed in the flanking cells of the *dlg1* RNAi embryos are less effective at mediating coupling between the actomyosin network and the cell boundary, and lead to “unclutched” contractions. **(B)** In control embryos, the flanking non-constricting cells (light blue) mediate mechanical coordination between the constriction domain (dark blue) and the non-constricting ectodermal tissue (white). This tissue-level coordination is important for an effective transition from apical constriction to invagination. When the mechanical integrity of the flanking cells is disrupted by 1) loss of apical-basal polarity (ie: *dlg1* RNAi embryo), 2) laser ablation, or 3) optogenetic downregulation of cortical actin, the flanking cells become overstretched (black cell outlines) by the constricting cells during apical constriction. Overstretching of the flanking cells impairs the mechanical coordination between the constricting cells and the surrounding ectodermal tissue, resulting in a delay in invagination.

While the mechanism by which Dlg1 regulates the properties of the flanking cells is not fully understood, our data suggest that the overstretched phenotype of the flanking cells is partially due to aberrant coupling of apical myosin contractions to the cell junctions. In both control and *dlg1* RNAi embryos, flanking cells accumulate a low level of apical myosin that undergoes contractions. We present evidence that apical myosin contractions in wildtype flanking cells temporarily stabilize apical area and prevent excessive apical stretching as the constricting cells apically constrict. Furthermore, we show that myosin pulses in wildtype flanking cells exhibit different levels of coupling to local changes in apical cell morphology. Pulses with stronger coupling are more effective at restraining cell stretching. Although the flanking cells in the *dlg1* RNAi embryos show a similar level of apical myosin accumulation and frequency of myosin coalescence, coupling between myosin pulses and local cell shape changes are reduced. This observation explains why myosin pulses are less effective at resisting cell stretching in the *dlg1* RNAi embryos.

The lower level of apical junctions observed in the flanking cells of the *dlg1* RNAi embryos can well explain the altered myosin behaviors and cell dynamics we have described. Weakening of adherens junctions is often associated with ineffective myosin-driven cell contractions. A previous study in *C. elegans* has shown that targeting a cadherin-catenin complex or the Rac pathway prevents effective ratcheting of cells during myosin contractions (Roh-Johnson et al., 2012). Disrupting the link between adherens junctions and the actomyosin network is also associated with ineffective myosin contractions. In *Drosophila* maternal/zygotic mutants for Canoe/Afadin, a linker between adherens junctions and the actin cytoskeleton, apical myosin accumulates and apical constriction initiates, but subsequently stalls (Sawyer et al., 2009). In this scenario, myosin contraction becomes uncoupled from cell shape changes, much like in the flanking cells of the *dlg1* RNAi embryos. We postulate that in wildtype flanking cells, the actomyosin fibers are strongly connected to the cell-cell junctions. This allows the apical myosin network to contract in a clutched/engaged manner and generate contractile forces on the cell-cell boundary to help resist pulling forces from the constriction domain, albeit temporarily. In the flanking cells of the *dlg1* RNAi embryos, attachment between the actomyosin network and the cell-cell junctions is impaired. In this case, the unclutched/disengaged actomyosin contractions are not sufficient to resist pulling forces from the constriction domain, and the flanking cells become hyper-stretched (Fig. 9A). Future experiments that manipulate adherens junctions in the flanking cells, without affecting the constricting cells, are necessary to test our working model.

We also investigated whether the mechanical properties of the flanking cells are altered in the *dlg1* RNAi embryos, which in principle could contribute to the overstretched phenotype of the flanking cells. Using magnetic tweezers, we applied pulling forces on a group of cells located in the ventral-lateral region of the cellular blastoderm. We found that compared to wildtype cells, cells in the *dlg1* RNAi embryos are less capable of restoring their original morphology after being stretched by magnetic tweezers. This suggests that the mechanical properties of these cells are affected by Dlg1 knockdown. It is possible that altered cell properties enhance the overstretched phenotype of the mutant flanking cells, as mutant cells were less capable of restoring their pre-stretched morphology when subjected to stretching. Although we were not able to specifically target the flanking cells in this experiment due to technical limitations, our data raises another potential factor that may account for the overstretched phenotype of the flanking cells.

Past studies of apical constriction mediated epithelial folding have mostly focused on the role of active forces generated in the constricting cells. The potential contribution of cells outside of the constriction domain has not been well studied. Our work indicates that the integrity of the flanking cells adjacent to the constriction domain can influence invagination by mediating coupling between the constricting cells and the ectodermal tissue. Furthermore, we demonstrate that this coupling requires the ability of the flanking cells to resist stretching from the constriction domain, which is dependent on clutched apical myosin contractions. In the future, it will be important to elucidate the actual mechanical contributions of non-constricting tissues during tissue folding and to further investigate how Dlg1 regulates mechanical properties at the cell and tissue level.

## Materials and Methods

### Fly stocks and genetics

E-cadherin-GFP (ubi-DE-cad–GFP) is described in (Morin et al., 2001). Sqh-GFP is described in (Royou et al., 2002). Sqh-mCherry is described in (Martin et al., 2009). Utr-Venus is a gift from the lab of A. Sokac (University of Illinois at Urbana-Champaign). UAS-Baz-GFP is a gift from the lab of Y.C. Wang (RIKEN). The following CPTI (Cambridge Protein Trap Insertion) stock was obtained from the Kyoto *Drosophila* Stock Center: Cno-YFP (115111). The following lines were obtained from the Bloomington *Drosophila* Stock Center: *dlg1* TRiP (36771, 33620), *dlg1*[2]/FM7a (36278), *dlg1*[5]/FM7a (36280), *scrib* TRiP (39073, 38199, 58085), and *lgl* TRiP (35773, 38989). The TRiP lines were crossed to a maternal GAL4 driver line, Maternal-Tubulin-Gal4 67.15 (“Mat67; Mat15”, (Hunter and Wieschaus, 2000)), for expression of shRNA during oogenesis. The following optogenetic lines were a gift from the De Renzis lab (EMBL): (1) w[*]; UASp-CIBN-pmGFP/CyO; Sb/TM3, (2) UASp-pmCIBN/FM6; Sb/TM6B, and (3) UASp-CRY2-OCRL/TM3.

To examine gastrulation defects in embryos with maternal knockdown of specific candidate genes via RNA interference (“RNAi embryos”), female flies from specific TRiP lines were crossed to Mat67 Sqh-mCherry; Mat15 E-cad-GFP/TM3 males to generate Mat67 Sqh-mCherry/TRiP; Mat15 E-cad-GFP/+ or Mat67 Sqh-mCherry/+; Mat15 E-cad-GFP/TRiP flies. The embryos derived from these flies were used for morphological analysis and analysis of adherens junction organization. To generate controls for the knockdown experiments, *y w f* females were crossed to Mat67 Sqh-mCherry; Mat15 E-cad-GFP/TM3 males to generate Mat67 Sqh-mCherry/+; Mat15 E-cad-GFP/+ flies. The embryos derived from these flies were used as control.

Similar crosses were made to examine the localization of specific proteins in RNAi embryos (F-actin, Cno/Afadin, and Baz/Par-3). The maternal GAL4 lines used for these studies were (1) Mat67 Sqh-mCherry; Mat15 Utr-Venus/TM3, (2) Mat67 Sqh-mCherry; Mat15 Cno-YFP/TM3, and (3) Mat67 Baz-GFP/Cyo; Mat15 Sqh-mCherry. In addition, embryos derived from Mat67 Sqh-mCherry/+; Mat15 Utr-Venus/*dlg1* TRiP flies were used for the magnetic tweezers experiments. Embryos derived from wildtype flies containing ubi-E-cadherin-GFP were used for laser ablation experiments.

To examine the actin phenotype in *dlg1* maternal mutant embryos, females from the *dlg1[5]*/FM7a; Utr-Venus Sqh-mCherry/TM3 stocks were crossed to *dlg1*[2]/Y males to generate *dlg1[2]*/*dlg1[5]*; Utr-Venus Sqh-mCherry/+ transheterozygous mutant flies. Embryos derived from these flies were imaged as *dlg1* maternal mutant embryos. *dlg1[2]*/FM7a; Utr-Venus Sqh-mCherry/+ heterozygous flies derived from the same cross were used to generate control embryos for this experiment.

For optogenetic disruption of F-actin, females from the UASp-pmCIBN/FM6; UASp-CRY2-OCRL/TM3 stock were crossed to Mat67 Sqh-mCherry; Mat15 E-cad-GFP/TM3 males to generate UASp-pmCIBN/+ or Y; Mat67 Sqh-mCherry/+; Mat15 E-cad-GFP/UASp-CRY2-OCRL flies. Embryos derived from these flies were used to determine the effect of stimulation on ventral furrow invagination. To confirm the loss of cortical actin upon stimulation, females from the UASp-pmCIBN/FM6; UASp-CRY2-OCRL/TM3 stock were crossed to Mat67 Sqh-mCherry; Mat15 Utr-Venus/TM3 males to generate UASp-pmCIBN/+ or Y; Mat67 Sqh-mCherry/+; Mat15 Utr-Venus/UASp-CRY2-OCRL flies. Embryos derived from these flies were used in this analysis.

### Immunostaining

Mat67 Sqh-mCherry/+; Mat15 E-cad-GFP/*dlg1* TRiP (*dlg1* RNAi) and Mat67 Sqh-mCherry/+; Mat15 E-cad-GFP/+ (control) flies were kept at 18°C and embryos were collected from overnight plates. Embryos were dechorionated with bleach for 1 minute, collected with a metal mesh, and rinsed extensively. Embryos were subsequently fixed with 10% paraformaldehyde for 1 hour. Following removal of the vitelline membrane, embryos were blocked with 10% bovine serum albumin (BSA) in PBS and 0.1% Tween 20. Primary antibody staining for Dlg1 (1:50; mouse, Developmental Studies Hybridoma Bank) and GFP (1:500; rabbit, EMD Millipore) was done in PBT (PBS/0.1% BSA/0.1% Tween 20) overnight at 4°C. Secondary antibody staining with Alexa Fluor 488 (1:500; anti-mouse; Invitrogen) and Alexa Fluor 561 (1:500; anti-rabbit; Invitrogen) was done for 1 hour at room temperature. Nuclei were stained with DAPI (1:10) for three minutes at room temperature. Embryos were mounted on glass slides containing Aqua Polymount (Polysciences). An A1 Nikon Confocal microscope with a 40× oil objective, a 405 nm laser, a 488 nm laser, and a 568 nm laser was used for imaging. Image size is 1024 pixels by 1024 pixels (0.21 μm/pixel).

### Confocal live imaging of ventral furrow formation and data analysis

All live imaging was performed at room temperature. Embryos were dechorionated in 3% bleach, rinsed thoroughly with water, transferred on a 35 mm MatTek glass-bottom dish (MatTek Corporation), and covered with water. In order to screen for invagination mutants, live imaging of control and RNAi embryos expressing a GFP tagged junctional/membrane marker, E-cadherin, and an mCherry tagged myosin marker, Spaghetti Squash, was performed using a Zeiss Axio Observer laser scanning confocal microscope (LSM 880), a 40×/1.3 numerical aperture oil-immersion objective, a 488 argon laser, and a 561 laser. A 1.57 × zoom was used. 9 – 11 confocal z-sections with a step size of 1 μm were acquired, with a temporal resolution ranging from 11 to 22.5 seconds. The image size was 512 pixels by 512 pixels, which corresponds to a lateral pixel size of 265 nm. The total imaged size is approximately 136 μm by 136 μm. The analyses of the confocal data are described below. All quantification and data plotting were performed using MATLAB (MathWorks), R, or Microsoft Excel unless otherwise stated.

#### (1) Analysis of tissue flow during ventral furrow formation

To measure tissue flow at the ventral surface of the embryo during ventral furrow formation (Figure 1), cartographic distortion due to the curvature of embryo surface was corrected by generating a flattened surface view as described by (Heemskerk and Streichan, 2015). Particle image velocimetry software (OpenPIV) (Taylor et al., 2010) was used to track the tissue movement towards the ventral midline from the surface view. A spacing/overlap of 32 pixels by 32 pixels and an interrogation window size of 32 pixels by 32 pixels was used. The average velocity of tissue movement towards the ventral midline (V_x_) was defined as the average velocity along the medial-lateral (ML) axis (the x direction under the described imaging setting) within a region 10 – 30 μm away from the ventral midline. Embryos were aligned in time based on the initial increase in V_x_ during apical constriction.

#### (2) Quantifying the rate of apical constriction

To measure the rate of apical constriction (Figure 1), individual constricting cells were segmented from the surface view of the movies using Embryo Development Geometry Explorer (EDGE), a MATLAB based image segmentation tool (Gelbart et al., 2012). E-cadherin-GFP was used as a membrane marker. EDGE detects membranes and fits individual cells into polygons. Manual corrections were carried out to ensure individual cells were segmented properly. The apical areas of individual cells within the constriction domain were traced 1 – 2 minutes before the onset of apical constriction and onwards until they disappeared from the surface view. The rate of constriction for individual cells was calculated as the rate of area reduction during the course of apical constriction.

#### (3) Quantifying the extent of flanking cell stretching

To determine the cell aspect ratio of the flanking cells (Figure 4), flanking cells at T_trans_ were segmented from the apical surface view using EDGE. The areas of the segmented cells were measured. To obtain the aspect ratio of the cell, each segmented cell was fit to an ellipse using the “fit_ellipse” function in MATLAB. Next, the intersection between the long or short axis of the fitted ellipse and the segmented cell boundary was determined and used as a measurement for the long or short axis of the cell, respectively. The aspect ratio of the cell is defined as the ratio between the long axis and short axis of the cell.

#### (4) Analysis of myosin pulses in the flanking cells

To analyze apical myosin in individual flanking cells (Figure 5), a z-plane close to the apical myosin signal was selected, and the E-cadherin-GFP signal was used to segment the cell outlines over time using EDGE. To select flanking cells, cells were first filtered based on their distance from the ventral midline at T_trans_ (16 – 40 μm). Flanking cell identity was then manually confirmed based on whether apical stretching was observed during apical constriction. To analyze apical myosin in the embryo, maximum projections of apical Sqh-mCherry signal within 5 µm from the apical surface were generated using the following approach in ImageJ. First, the z-slice with the brightest apical myosin signal at T_trans_ was determined (“Z_a_”). Next, a sub-stack including Z_a_ and all z-slices 5 µm below Z_a_ was generated. Next, a gaussian blur filter with a Sigma (Radius) of 1 (pixel) was applied to each z-slice in the sub-stack to reduce noise. Finally, a maximum intensity projection was generated from the gaussian blurred sub-stack. Myosin intensities between different embryos were normalized based on the background apical myosin intensity at a reference timepoint (T_ref_) that was within one minute before T_trans_. Background myosin intensity (“Myo_BG_”) was defined as the myosin intensity in regions where no myosin pulses were detected at T_ref_ and was determined as follows. For each embryo, the mean pixel intensity of apical myosin for each segmented cell was measured and plotted as a function of the position of the cell relative to the ventral midline at T_trans_. The myosin intensities lowered quickly from 0 μm to 15 μm and reached the floor at 20 – 30 μm. Myo_BG_ was defined as the myosin intensity at the floor. A scaling factor (sf) was then calculated by dividing Myo_BG_ by the mean myosin background intensity of all embryos analyzed. For each embryo, the normalized apical myosin intensity (Myo_NOR_) for a single segmented cell was calculated as:

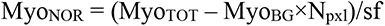

where Myo_TOT_ is the total myosin intensity within the apical domain of the cell and N_pxl_ is the size of the apical domain in pixels. For each segmented flanking cell, the normalized apical myosin intensities were plotted over time as a heatmap to reveal cycles of apical myosin accumulation and disappearance.

To analyze myosin pulses in the flanking cells, individual myosin coalescence events were manually identified. For each event, the timepoint when apical myosin coalesced into a focused, high-intensity punctum was defined as the pulse peak. A 96-second time interval centered around the pulse peak, which we referred to as a myosin “pulse”, was used for analysis of changes in apical cell area, apical cell length, and apical myosin intensity. The 96-second time interval was chosen in order to maximize coverage of the rising and falling phases of a myosin coalescence event, and to minimize the overlap between two consecutive myosin coalescence events. To compare cell behaviors during pulses and time intervals between pulses, 96-second intervals centered around timepoints showing local minimums of myosin intensity were selected as “off-pulses” and analyzed. Off-pulses served as controls for pulses. To analyze the position of the intensity peak of apical myosin, the weighted centroid of apical Sqh-mCherry signal within each flanking cell was determined at the pulse peak time and expressed as a percentage relative to the leading and lagging edge of the cell (the sides of the cell closest to and furthest away from the ventral midline, respectively). The leading edge and the lagging edge were defined as 0% and 100%, respectively. To analyze the coupling between myosin coalescence and local cell deformation, the change in the width of the cell along the anterior-posterior direction at the position where myosin coalescence occurred was manually measured in ImageJ as a “Local contraction index” to represent local cell shape change associated with the myosin coalescence.

### Imaging and analysis of the localization of subapical landmarks

In order to examine the localization of subapical components during polarity establishment, live imaging of control and *dlg1* mutant embryos expressing Canoe-YFP, Bazooka-GFP, E-cadherin-GFP, (Fig. S3) or Utrophin-Venus (Fig. S4) was performed with an Olympus FVMPE-RS multiphoton microscope, a 25×/1.05 numerical aperture water-immersion objective, and a 920 nm pulsed laser. A 1× zoom was used. Single plane images (1024 pixels by 1024 pixels, or 509 µm by 509 µm) at the midsagittal plane of the embryo were acquired in 1-minute intervals until germband extension. The lateral pixel size is 0.50 μm. Embryos were mounted lateral side up such that both dorsal and ventral sides of the embryo were captured when imaged at the midsagittal plane. In order to quantify actin distribution along the lateral membrane of the non-constricting ectodermal cells, the image frame corresponding to the onset of ventral furrow formation was selected, and a region of interest (251 pixels by 93 pixels, 125 μm by 46 μm) covering approximately 24 epithelial cells at the dorsal region of the embryo was used for quantification. The positions of the lateral membranes were manually segmented by drawing a line along each lateral cortical F-actin signal. Mean Utr-Venus intensity was integrated along each segmented lateral membrane with a width of five pixels (2.5 μm). The measured intensity was subsequently normalized between embryos based on the mean signal intensity within the epithelial layer at the dorsal side of the embryo. Between 14 and 24 lateral membranes were measured for each embryo and averaged to determine the apical-basal distribution of cortical actin. Finally, the apical-basal distribution of cortical actin signal in embryos with the same maternal genotype were averaged to generate the mean actin distribution.

### Multiphoton-based deep tissue live-imaging and data analysis

Deep tissue live-imaging of ventral furrow formation in control and *dlg1* RNAi embryos was performed with an Olympus FVMPE-RS multiphoton microscope, a 25×/1.05 numerical aperture water-immersion objective, and a 920 nm pulsed laser. A 3× zoom was used. Embryos were mounted ventral side up. For different batches of experiments, two similar acquisition conditions were used. In the first condition, stacks of 51 images taken at 2-μm steps were acquired during ventral furrow formation in 2-minute intervals with a resonant scanner. A region of interest (512 pixels by 256 pixels, 170 μm by 85 μm) that encompassed the mid-region of the embryo (around 50% of egg length) was imaged. In the second condition, stacks of 100 images taken at 1-μm steps were acquired in 2-minute intervals with a Galvano scanner. A smaller region of interest, 512 pixels by 128 pixels (170 μm by 42 μm), was imaged. In both imaging conditions, laser power was increased linearly over the span of 100 μm in order to compensate for the loss of signal due to light scattering. The analyses of the multiphoton data are described below.

#### (1) Measuring the rate of ventral furrow invagination

To measure the rate of ventral furrow invagination (Fig. 2A-F), cross section views encompassing the entire depth of the ventral furrow were generated in ImageJ through re-slicing, followed by an average projection of 20 slices. Embryos were aligned by time based on the onset of apical constriction. The rate of furrow ingression was measured by manually tracking the distance between the vitelline membrane and the apex of the ventral most cell (D) over time during ventral furrow formation.

#### (2) Analyzing the 3D morphology of the ventral furrow at T_trans_

The following analyses were performed to evaluate the morphology of the intermediate furrow at T_trans_ (defined as 8 minutes after the onset of apical constriction for multiphoton movies with a 2-minute resolution) (Fig. 2G-J). First, the apical and basal arcs of the ventral furrow at T_trans_ were manually outlined from the cross-section view in order to quantify the apical and basal arc length. Second, the furrow thickness was measured by determining the distance between the apical and basal membranes of the cells closest to the ventral midline. Finally, a single row of cells encompassing the constriction domain and the flanking cells were segmented in 3D using EDGE and the surfaces of the segmented cells were plotted in 3D for comparison.

#### (3) Analyzing mesodermal and ectodermal cell movement during ventral furrow formation

To analyze the ventrally directed movement of the non-constricting cells towards the ventral midline during ventral furrow formation (Fig. 6), a flattened ventral surface view was generated from each multiphoton movie using a custom MATLAB script. A row of ∼12 cells on one side of the ventral midline that included ∼ 6 constricting cells, 3 flanking cells, and ∼ 3 ectodermal cells was segmented from the surface view using EDGE and manually tracked over time. Quantities such as apical cell area, the medial-lateral (M-L) position of the cell apex, and the distance traveled to the midline were extracted from the segmented cells. The measurements were further interpolated based on the initial M-L position of the cells at the onset of apical constriction and subsequently averaged between embryos from the same genotype. The velocity of cell movement between 0 – 8 minutes was calculated based on the measurement of cell position over time.

### Magnetic Tweezers

Embryos were dechorionated and mounted ventral side down on a 24 mm × 50 mm 1.5 coverslip containing a thin layer of glue on the surface. Once mounted, the embryos were dried in a desiccator for 12 minutes. After desiccation, embryos were covered with a 3:1 mixture of 700/27 halocarbon oil. Yellow fluorescent carboxyl magnetic particles that are 0.5 μm in diameter (Spherotech) were diluted in water (1:10) and injected into the periplasm of the embryos during cellularization using a FemtoJet express microinjector (Eppendorf). The embryos used in this experiment expressed Utr-Venus and Sqh-mCherry which allowed us to monitor cell shape and actomyosin dynamics. A home-built electromagnet was used to pull on the magnetic beads.

Before each experiment, a z-stack spanning approximately 1/3 of the embryo’s thickness, taken at 1-μm steps, was acquired to record the distribution of magnetic beads within the embryo. Simultaneous dual-color imaging was performed with an inverted Nikon spinning disk confocal Ti microscope, a 488 nm laser, a 561 nm laser, and a 40× oil-immersion objective. Images are 2048 pixels by 2048 pixels (340 μm by 340 μm). The lateral pixel size is 166 nm. The total intensity of beads within each embryo was determined by subtracting the background levels of fluorescence from the total image intensity. Since the magnetic beads were much brighter than Utr-Venus, the presence of Utr-Venus in the embryos did not noticeably affect the quantification of the intensity of the beads.

To capture the response of the magnetic beads to pulling from the electromagnet, simultaneous dual-color imaging was performed with a Nikon spinning disk confocal Ti microscope, a 488 nm laser, a 561 nm laser, and a 40× oil-immersion objective. Images 2048 pixels by 2048 pixels (340 μm by 340 μm) in size were acquired at a single focal plane within the periplasm every 0.43 seconds. After image acquisition, the displacement of the magnetic beads along the direction of pulling was manually tracked over time. Three clusters of magnetic beads near the center of the bead distribution were chosen for manual tracking. The position of a group of cells away from the clusters of magnetic beads was measured as a control. In some instances, embryos were slightly rotated within the eggshell upon pulling. Embryo rotation was detected when cells located far away from the beads (the “control cells”) moved in the direction of pulling as a cohort without showing obvious local cell shape changes. To account for this global movement, we generated rotation-corrected displacement curves by subtracting the displacement of the control cells from that of the magnetic beads. For each embryo, the average bead displacement curve was generated from the displacement curves of three clusters of manually tracked beads. The displacement curves include displacement of the beads during the resting phase (60 seconds), the pulling phase (30 seconds), and the recovery phase (∼ 120 seconds). The maximum displacement of the beads was set as the total bead displacement during the entire pulling phase. The percent recovery of the beads was calculated by dividing the net displacement of beads during the first 100 seconds of the recovery phase by the total displacement of the beads during the pulling phase.

### Laser Ablation

Embryos expressing E-cadherin-GFP were prepared in the same way as for regular live imaging described above. Embryos were mounted ventral side up and imaged with an Olympus FVMPE-RS multiphoton microscope, a 25×/1.05 numerical aperture water-immersion objective, a 920 nm pulsed laser, and a Galvano scanner. A 3× zoom was used. For each laser ablation experiment, an embryo at the mid-phase of apical constriction was selected, and the following three steps were repeated for 30 cycles. In the first step, z-stacks of 21 images taken at 2-μm steps were acquired for two timepoints in 8-second intervals, for a total of 16-seconds. Images encompass the apical region of the embryo and are 512 pixels by 128 pixels (170 μm by 42 μm) in size. In the second step, laser ablation of one strip of flanking cells 50 pixels by 250 pixels in size (17 μm along the medial-lateral axis and 83 μm along the anterior-posterior axis) was performed on both sides of the constriction domain. Laser ablation was done for 2-seconds on the apical-most domain of the flanking cells and spanned a depth of 2-μm. The third step was similar to the first step, except that six timepoints were acquired for a total of 48-seconds. The total length of duration for all three cycles was 68-seconds, and for 30 iterations it took approximately 35-minutes. In this protocol, the flanking cells were subjected to laser ablation for 2 seconds every 68 seconds. We found that repeating laser ablation is necessary to prevent active shrinking of the cut apical domain, which typically happened ∼ 1 minute after cutting. A similar protocol was used for the control experiments, except that a much lower laser power was used in step 2, such that only photobleaching occurred and there was no visible recoil indicative of laser ablation. After image acquisition, cross-section views were generated in ImageJ by re-slicing, followed by an average intensity projection of 20 slices. In addition, ventral surface views were generated from the original image stacks using a custom MATLAB script. The rate of ventral furrow invagination was manually tracked from the cross-section view by measuring the invagination depth, D, over time after laser ablation. Embryos were aligned in time by T_trans_ in order to compare the rate of invagination.

### Optogenetics

Embryos were prepared in the same way as for regular live imaging, except that the preparation was performed either in the dark or under red light. Single color imaging was performed with an Olympus FVMPE-RS multiphoton microscope, a 25×/1.05 numerical aperture water-immersion objective, a 1040 nm laser line, and a resonant scanner. A 3× zoom was used. Stacks of 101 images taken at 1-μm steps were acquired in 1-minute intervals before and after optogenetic stimulation. Images encompassed the mid-region of the embryo (around 50% of the egg length) and were 512 pixels by 512 pixels (170 μm by 170 μm) in size. Optogenetic stimulation of a strip of non-constricting cells (170 μm along the anterior-posterior axis and ∼ 30 μm along the medial-lateral axis) was performed during apical constriction through continuous excitation with a 920 nm pulsed laser for approximately 2 minutes with Galvano scanning. The stimulated region included the flanking cells and ∼ 1 – 2 rows of ectodermal cells at one side of the constriction domain. Embryos with an identical maternal genotype but imaged without stimulation were used as control for this experiment. Cross section views were generated in ImageJ. Ventral surface views were generated from the original image stacks using a custom MATLAB script. The alignment of time and the rate of ventral furrow ingression was determined in the same way as in the laser ablation experiments.

### Statistical Analysis

Sample sizes for the presented data and methods for statistical comparisons can be found in figure legends. *p* values were calculated using MATLAB ttest2 or ranksum function.

## Author contributions

Conceptualization: B.H., M.A.F.; Methodology: B.H., M.A.F.; Software: B.H., M.A.F.; Validation: B.H., M.A.F.; Formal analysis: B.H., M.A.F.; Investigation: M.A.F.; Resources: B.H.; Data Curation: M.A.F.; Writing – original draft preparation: M.A.F.; Writing-review and editing: B.H., M.A.F.; Visualization: B.H., M.A.F.; Supervision: B.H.; Project administration: B.H.; Funding acquisition: B.H.

## Acknowledgements

We thank A. Lavanway for critical help with imaging and research support in general. We thank B. Robertson for help with building the electromagnet used in our experiments. We thank B. J. Gourgeot and K. P. Royce for help with pilot data analysis. We thank S. De Renzis and Y.C. Wang for sharing reagents. We thank E. Griffin and J. Moseley for their valuable feedback on our manuscript. We thank M. Peifer and T. Bonello for helpful discussion. We thank the B.H. lab members for constructive comments and discussions. We thank the Bloomington Drosophila Stock Center and the Developmental Studies Hybridoma Bank for providing reagents used in this work. This research is supported by NIGMS ESI-MIRA R35GM128745 and American Cancer Society Research Grant #IRG-16-191-33 to B.H, and the GAANN fellowship and the Ryan fellowship to M.A.F.

## References

1. Barrett, K., Leptin, M. and Settleman, J. (1997). The Rho GTPase and a putative RhoGEF mediate a signaling pathway for the cell shape changes in Drosophila gastrulation. Cell 91, 905–915.

2. Bilder, D., Li, M. and Perrimon, N. (2000). Cooperative regulation of cell polarity and growth by Drosophila tumor suppressors. Science 289, 113–116.

3. Bonello, T. T., Choi, W. and Peifer, M. (2019). Scribble and Discs-large direct initial assembly and positioning of adherens junctions during the establishment of apical-basal polarity. Development 146, dev180976.

4. Costa, M., Wilson, E. T. and Wieschaus, E. (1994). A putative cell signal encoded by the folded gastrulation gene coordinates cell shape changes during Drosophila gastrulation. Cell 76, 1075–1089.

5. Dawes-Hoang, R. E., Parmar, K. M., Christiansen, A. E., Phelps, C. B., Brand, A. H. and Wieschaus, E. F. (2005). folded gastrulation, cell shape change and the control of myosin localization. Development 132, 4165–4178.

6. Denk-Lobnig, M., Totz, J. F., Heer, N. C., Dunkel, J. and Martin, A. C. (2021). Combinatorial patterns of graded RhoA activation and uniform F-actin depletion promote tissue curvature. Development 148, dev199232.

7. Fernandez-Gonzalez, R. and Zallen, J. A. (2011). Oscillatory behaviors and hierarchical assembly of contractile structures in intercalating cells. Phys Biol 8, 045005.

8. Fuse, N., Yu, F. and Hirose, S. (2013). Gprk2 adjusts Fog signaling to organize cell movements in Drosophila gastrulation. Development 140, 4246–4255.

9. Gelbart, M. A., He, B., Martin, A. C., Thiberge, S. Y., Wieschaus, E. F. and Kaschube, M. (2012). Volume conservation principle involved in cell lengthening and nucleus movement during tissue morphogenesis. Proc Natl Acad Sci U S A 109, 19298–19303.

10. Gracia, M., Theis, S., Proag, A., Gay, G., Benassayag, C. and Suzanne, M. (2019). Mechanical impact of epithelial-mesenchymal transition on epithelial morphogenesis in Drosophila. Nat Commun 10, 2951.

11. Guglielmi, G., Barry, J. D., Huber, W. and De Renzis, S. (2015). An Optogenetic Method to Modulate Cell Contractility during Tissue Morphogenesis. Dev Cell 35, 646–660.

12. Häcker, U. and Perrimon, N. (1998). DRhoGEF2 encodes a member of the Dbl family of oncogenes and controls cell shape changes during gastrulation in Drosophila. Genes Dev 12, 274–284.

13. Heemskerk, I. and Streichan, S. J. (2015). Tissue cartography: compressing bio-image data by dimensional reduction. Nat Methods 12, 1139–1142.

14. Hunter, C. and Wieschaus, E. (2000). Regulated expression of nullo is required for the formation of distinct apical and basal adherens junctions in the Drosophila blastoderm. J Cell Biol 150, 391–401.

15. John, A. and Rauzi, M. (2021). A two-tier junctional mechanism drives simultaneous tissue folding and extension. Dev Cell 56, 1469–1483.e5.

16. Kerridge, S., Munjal, A., Philippe, J.-M., Jha, A., de las Bayonas, A. G., Saurin, A. J. and Lecuit, T. (2016). Modular activation of Rho1 by GPCR signalling imparts polarized myosin II activation during morphogenesis. Nat Cell Biol 18, 261–270.

17. Kim, H. Y., Varner, V. D. and Nelson, C. M. (2013). Apical constriction initiates new bud formation during monopodial branching of the embryonic chicken lung. Development 140, 3146–3155.

18. Kölsch, V., Seher, T., Fernandez-Ballester, G. J., Serrano, L. and Leptin, M. (2007). Control of Drosophila gastrulation by apical localization of adherens junctions and RhoGEF2. Science 315, 384–386.

19. Krueger, D., Tardivo, P., Nguyen, C. and De Renzis, S. (2018). Downregulation of basal myosin-II is required for cell shape changes and tissue invagination. EMBO J 37, e100170.

20. Leptin, M. (1991). twist and snail as positive and negative regulators during Drosophila mesoderm development. Genes Dev 5, 1568–1576.

21. Leptin, M. (1999). Gastrulation in Drosophila: the logic and the cellular mechanisms. EMBO J 18, 3187–3192.

22. Leptin, M. and Grunewald, B. (1990). Cell shape changes during gastrulation in Drosophila. Development 110, 73–84.

23. Liu, H., Yu, X., Li, K., Klejnot, J., Yang, H., Lisiero, D. and Lin, C. (2008). Photoexcited CRY2 interacts with CIB1 to regulate transcription and floral initiation in Arabidopsis. Science 322, 1535–1539.

24. Manning, A. J., Peters, K. A., Peifer, M. and Rogers, S. L. (2013). Regulation of epithelial morphogenesis by the G protein-coupled receptor mist and its ligand fog. Sci Signal 6, ra98.

25. Martin, A. C. and Goldstein, B. (2014). Apical constriction: themes and variations on a cellular mechanism driving morphogenesis. Development 141, 1987–1998.

26. Martin, A. C., Kaschube, M. and Wieschaus, E. F. (2009). Pulsed contractions of an actin-myosin network drive apical constriction. Nature 457, 495–499.

27. Martin, A. C., Gelbart, M., Fernandez-Gonzalez, R., Kaschube, M. and Wieschaus, E. F. (2010). Integration of contractile forces during tissue invagination. J Cell Biol 188, 735–749.

28. Mason, F. M., Tworoger, M. and Martin, A. C. (2013). Apical domain polarization localizes actin-myosin activity to drive ratchet-like apical constriction. Nat Cell Biol 15, 926–936.

29. Morin, X., Daneman, R., Zavortink, M. and Chia, W. (2001). A protein trap strategy to detect GFP-tagged proteins expressed from their endogenous loci in Drosophila. Proc Natl Acad Sci U S A 98, 15050–15055.

30. Munjal, A. and Lecuit, T. (2014). Actomyosin networks and tissue morphogenesis. Development 141, 1789–1793.

31. Ni, J.-Q., Zhou, R., Czech, B., Liu, L.-P., Holderbaum, L., Yang-Zhou, D., Shim, H.-S., Tao, R., Handler, D., Karpowicz, P., et al. (2011). A genome-scale shRNA resource for transgenic RNAi in Drosophila. Nat Methods 8, 405–407.

32. Nikolaidou, K. K. and Barrett, K. (2004). A Rho GTPase signaling pathway is used reiteratively in epithelial folding and potentially selects the outcome of Rho activation. Curr Biol 14, 1822–1826.

33. Nishimura, T., Honda, H. and Takeichi, M. (2012). Planar cell polarity links axes of spatial dynamics in neural-tube closure. Cell 149, 1084–1097.

34. Parks, S. and Wieschaus, E. (1991). The Drosophila gastrulation gene concertina encodes a G alpha-like protein. Cell 64, 447–458.

35. Perez-Mockus, G., Mazouni, K., Roca, V., Corradi, G., Conte, V. and Schweisguth, F. (2017). Spatial regulation of contractility by Neuralized and Bearded during furrow invagination in Drosophila. Nat Commun 8, 1594.

36. Perrimon, N. (1988). The maternal effect of lethal(1)discs-large-1: a recessive oncogene of Drosophila melanogaster. Dev Biol 127, 392–407.

37. Polyakov, O., He, B., Swan, M., Shaevitz, J. W., Kaschube, M. and Wieschaus, E. (2014). Passive mechanical forces control cell-shape change during Drosophila ventral furrow formation. Biophys J 107, 998–1010.

38. Rauzi, M., Lenne, P.-F. and Lecuit, T. (2010). Planar polarized actomyosin contractile flows control epithelial junction remodelling. Nature 468, 1110–1114.

39. Rauzi, M., Krzic, U., Saunders, T. E., Krajnc, M., Ziherl, P., Hufnagel, L. and Leptin, M. (2015). Embryo-scale tissue mechanics during Drosophila gastrulation movements. Nat Commun 6, 8677.

40. Roh-Johnson, M., Shemer, G., Higgins, C. D., McClellan, J. H., Werts, A. D., Tulu, U. S., Gao, L., Betzig, E., Kiehart, D. P. and Goldstein, B. (2012). Triggering a cell shape change by exploiting preexisting actomyosin contractions. Science 335, 1232–1235.

41. Royou, A., Sullivan, W. and Karess, R. (2002). Cortical recruitment of nonmuscle myosin II in early syncytial Drosophila embryos: its role in nuclear axial expansion and its regulation by Cdc2 activity. J Cell Biol 158, 127–137.

42. Sawyer, J. K., Harris, N. J., Slep, K. C., Gaul, U. and Peifer, M. (2009). The Drosophila afadin homologue Canoe regulates linkage of the actin cytoskeleton to adherens junctions during apical constriction. J Cell Biol 186, 57–73.

43. Sawyer, J. M., Harrell, J. R., Shemer, G., Sullivan-Brown, J., Roh-Johnson, M. and Goldstein, B. (2010). Apical constriction: a cell shape change that can drive morphogenesis. Dev Biol 341, 5–19.

44. Sawyer, J. K., Choi, W., Jung, K.-C., He, L., Harris, N. J. and Peifer, M. (2011). A contractile actomyosin network linked to adherens junctions by Canoe/afadin helps drive convergent extension. Mol Biol Cell 22, 2491–2508.

45. Sherrard, K., Robin, F., Lemaire, P. and Munro, E. (2010). Sequential activation of apical and basolateral contractility drives ascidian endoderm invagination. Curr Biol 20, 1499–1510.

46. Sweeton, D., Parks, S., Costa, M. and Wieschaus, E. (1991). Gastrulation in Drosophila: the formation of the ventral furrow and posterior midgut invaginations. Development 112, 775–789.

47. Tanentzapf, G. and Tepass, U. (2003). Interactions between the crumbs, lethal giant larvae and bazooka pathways in epithelial polarization. Nat Cell Biol 5, 46–52.

48. Taylor, Z. J., Gurka, R., Kopp, G. A. and Liberzon, A. (2010). Long-Duration Time-Resolved PIV to Study Unsteady Aerodynamics. IEEE Trans. Instrum. Meas. 59, 3262– 3269.

49. Woods, D. F., Hough, C., Peel, D., Callaini, G. and Bryant, P. J. (1996). Dlg protein is required for junction structure, cell polarity, and proliferation control in Drosophila epithelia. J Cell Biol 134, 1469–1482.

50. Zhang, X., Jefferson, A. B., Auethavekiat, V. and Majerus, P. W. (1995). The protein deficient in Lowe syndrome is a phosphatidylinositol-4,5-bisphosphate 5-phosphatase. Proc Natl Acad Sci U S A 92, 4853–4856.

