## Supplemental Materials for "Cell polarity determinant Dlg1 facilitates epithelial invagination by promoting tissue-scale mechanical coordination"

#### Supplemental Figures 1 – 9

#### Supplemental Movies 1 – 6

#### Supplemental Materials

#### Figure S1

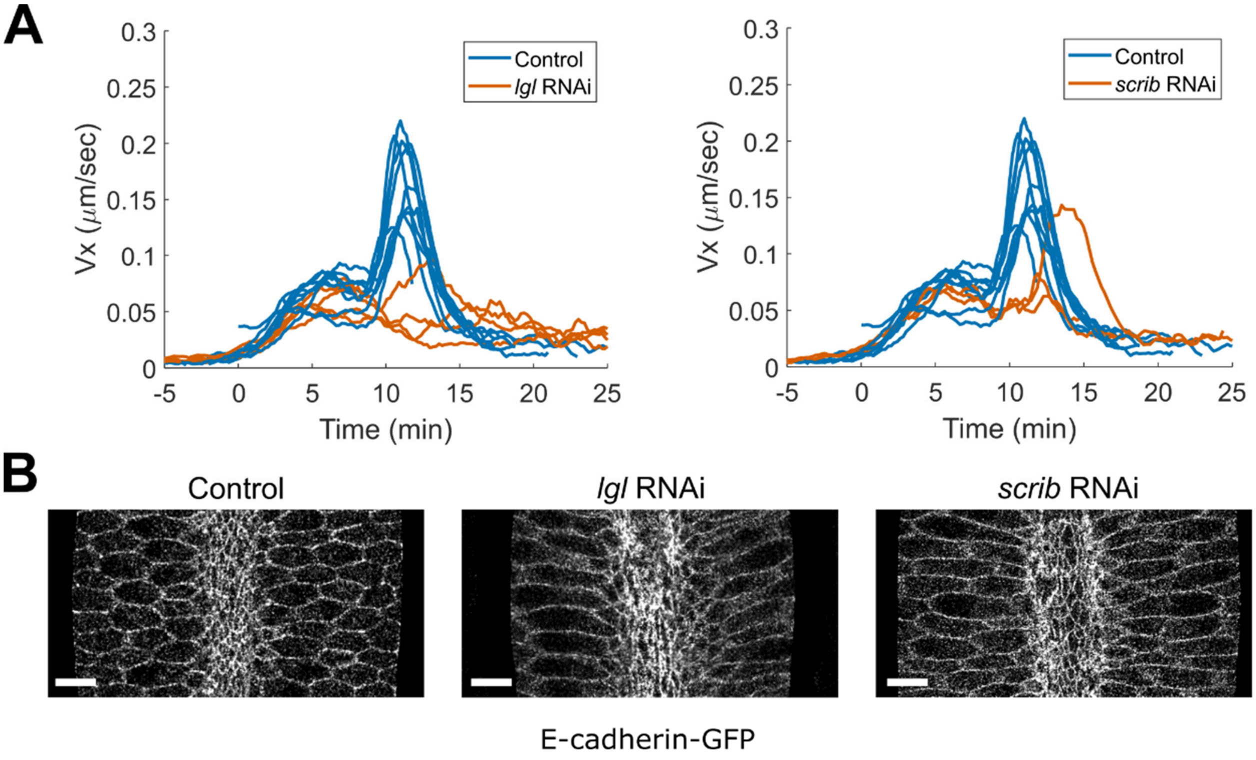


**Figure S1. *lgl* and *scrib* RNAi embryos show similar defects in ventral furrow invagination as *dlg1* RNAi embryos**

**(A)** PIV analysis of tissue movement during ventral furrow formation in *scrib* and *lgl* RNAi embryos. The average velocity of tissue movement 10 – 30 μm away from the ventral midline ($V_x$) was plotted over time. The same control embryos shown in Figure 1F were replotted for comparison here. **(B)** The flanking cells in *scrib* and *lgl* RNAi embryos become overstretched during ventral furrow formation. The cell membrane was visualized using E-cadherin-GFP. Scale bars: 10 μm.

**Figure S2**

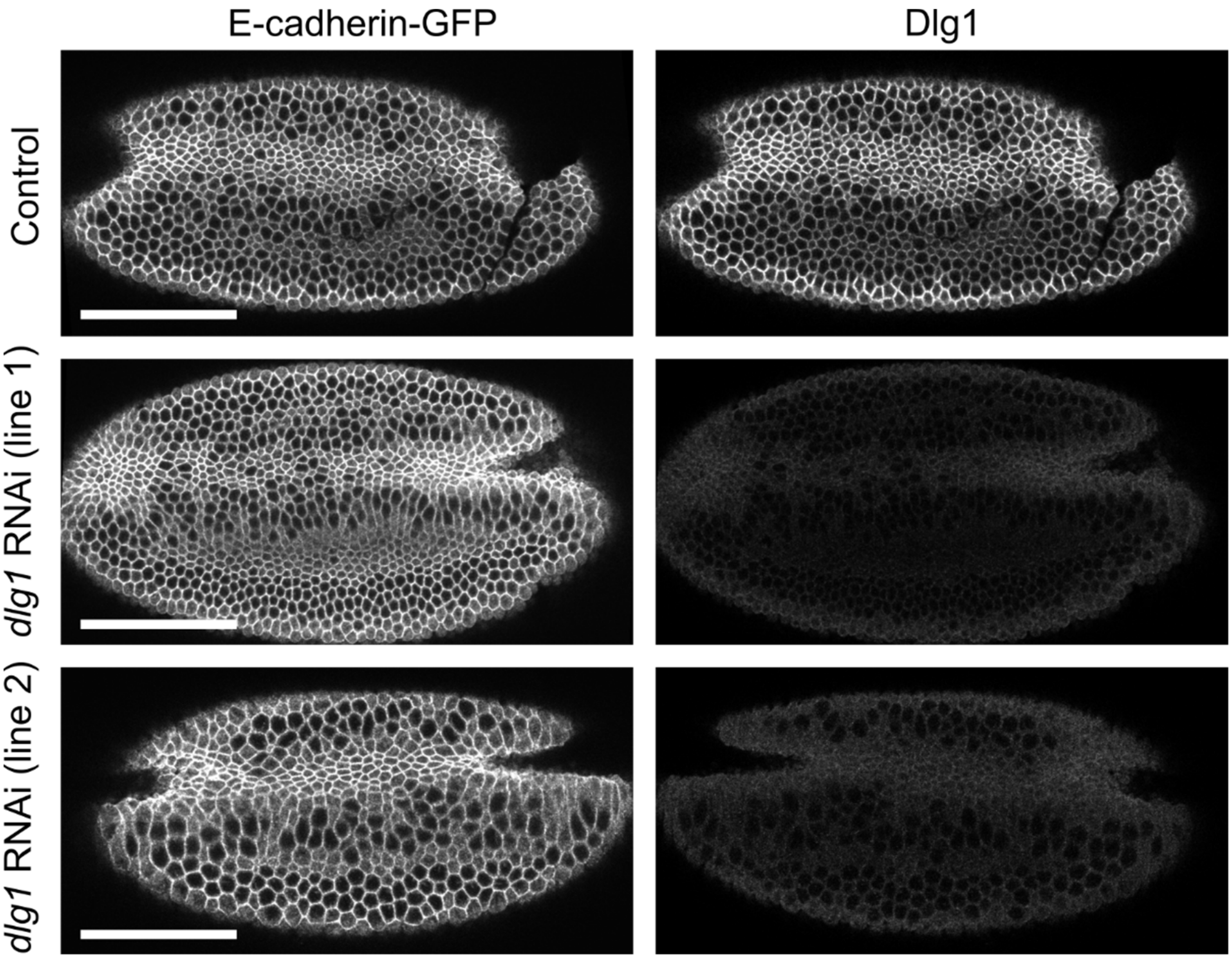


**Figure S2. Effective knockdown of Dlg1 in early embryos by transgenic RNAi**

Immunostaining of control and *dlg1* RNAi embryos expressing E-cadherin-GFP with anti-GFP and anti-Dlg1 antibodies. Shown are embryos undergoing apical constriction. Dlg1 signal is barely detectable in *dlg1* RNAi embryos. Scale bar: 50 µm.

#### Figure S3

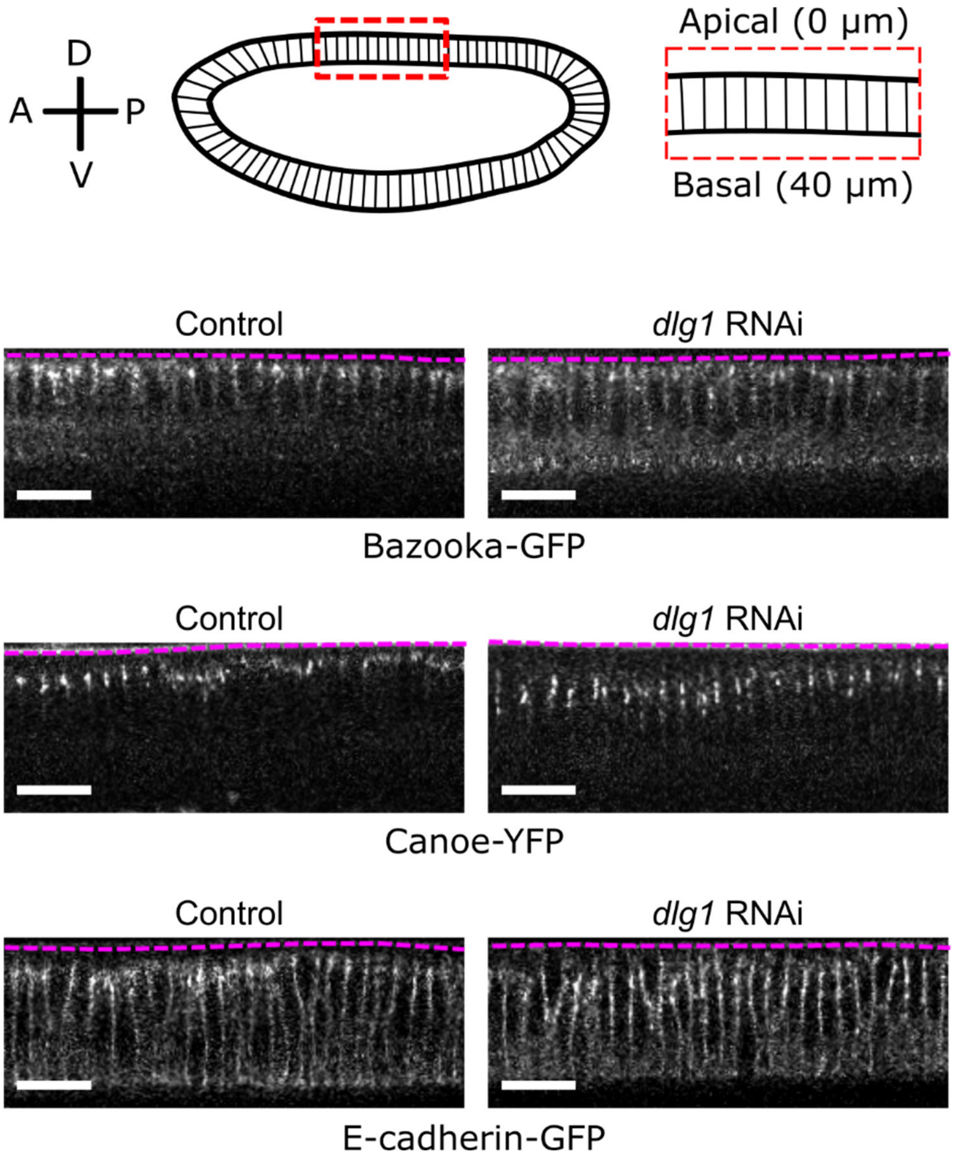


**Figure S3. Bazooka/Par-3, E-cadherin, and Canoe/Afadin distribution in dorsal ectodermal cells of control and *dlg1* RNAi embryos at the end of cellularization**

The localization pattern of Bazooka/Par-3 (Baz), Canoe/Afadin (Cno), and E-cadherin along the lateral membrane is consistent across multiple control (n = 3, 5, 17 for i, ii, and iii, respectively) and *dlg1* RNAi (n = 3, 10, 9 for i, ii, and iii, respectively) embryos. *dlg1* RNAi embryos exhibit a basal shift/spread of Baz/Par-3, Cno/Afadin, and E-cadherin signal. Scale bars: 20 μm.

#### Figure S4

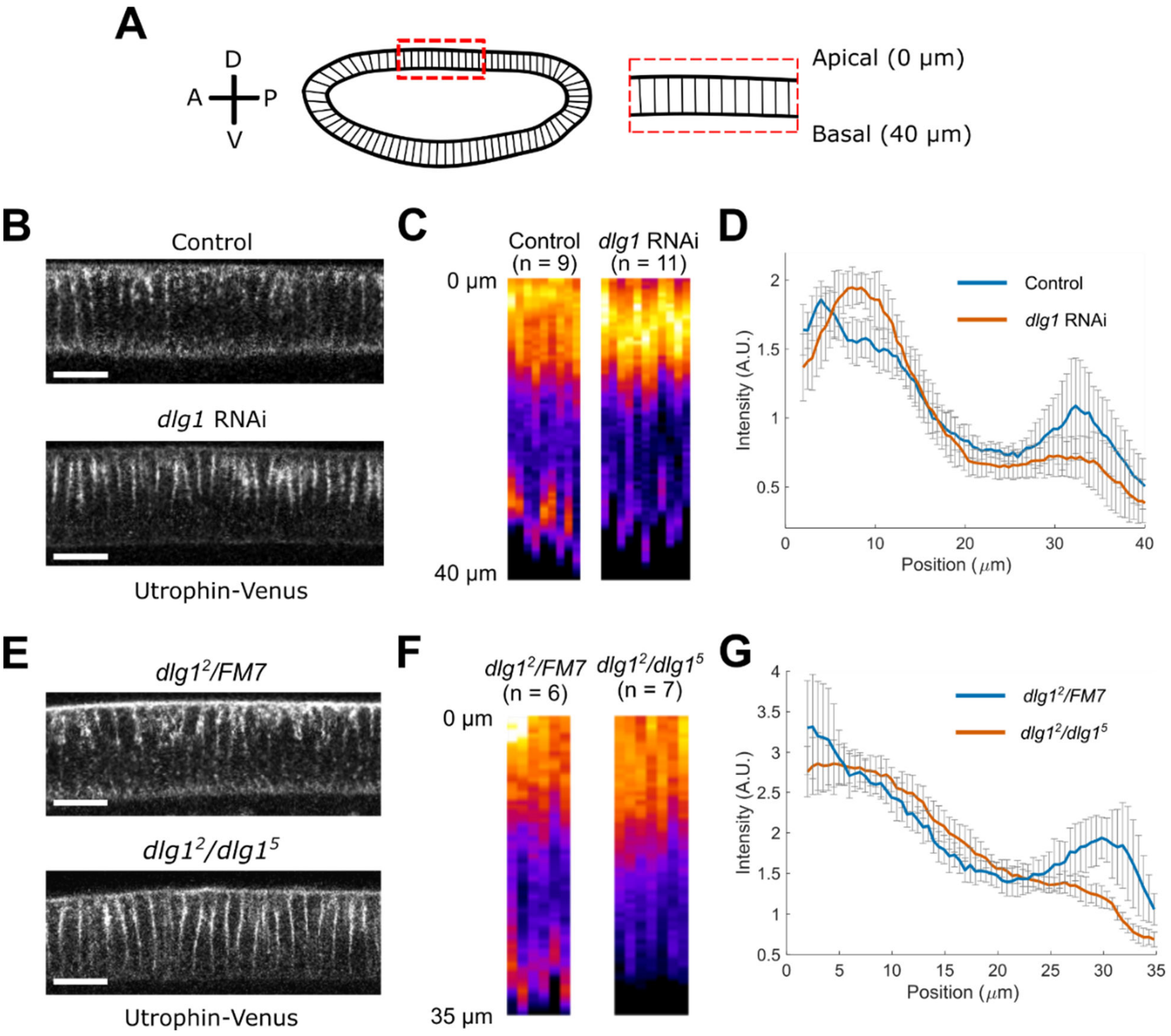


**Figure S4. Actin distribution along the lateral membrane of dorsal ectodermal cells is altered in *dlg1* mutant embryos**

**(A)** Top panel: Embryos were imaged laterally with the anterior side to the left and the dorsal side facing up, as illustrated in the cartoon. Intensity measurements were performed on a subset of the dorsal non-constricting ectodermal cells (red box). **(B, E)** Representative image highlighting F-actin distribution along the lateral membrane of the dorsal non-constricting ectodermal cells in a control and *dlg1* RNAi embryo (B), and in a *dlg1*$^{2}$/+ and *dlg1*$^{2}$/*dlg1*$^{5}$ embryo (E) at the onset of gastrulation (T = 0:00 (mm:ss)). F-actin was visualized using Utr-Venus. Scale bars: 20 µm. **(C, F)** Utr-Venus intensity along the lateral membrane of individual control (n = 9) and *dlg1* RNAi (n = 11) embryos (C), and individual *dlg1*$^{2}$/+ (n = 6) and *dlg1*$^{2}$/*dlg1*$^{5}$ (n = 7) embryos (F) at T = 0:00 (mm:ss). The heatmaps are oriented in the apical to basal direction from top to bottom. **(D, G)** Average actin intensity along the lateral membrane of dorsal non-constricting ectodermal cells for multiple control (n = 9) and *dlg1* RNAi (n = 11) embryos (D), and multiple *dlg1*$^{2}$/+ (n = 6) and *dlg1*$^{2}$/*dlg1*$^{5}$ (n = 7) embryos (G). The position is oriented in the apical to basal direction. Error bar represents the standard deviation.

### Figure S5

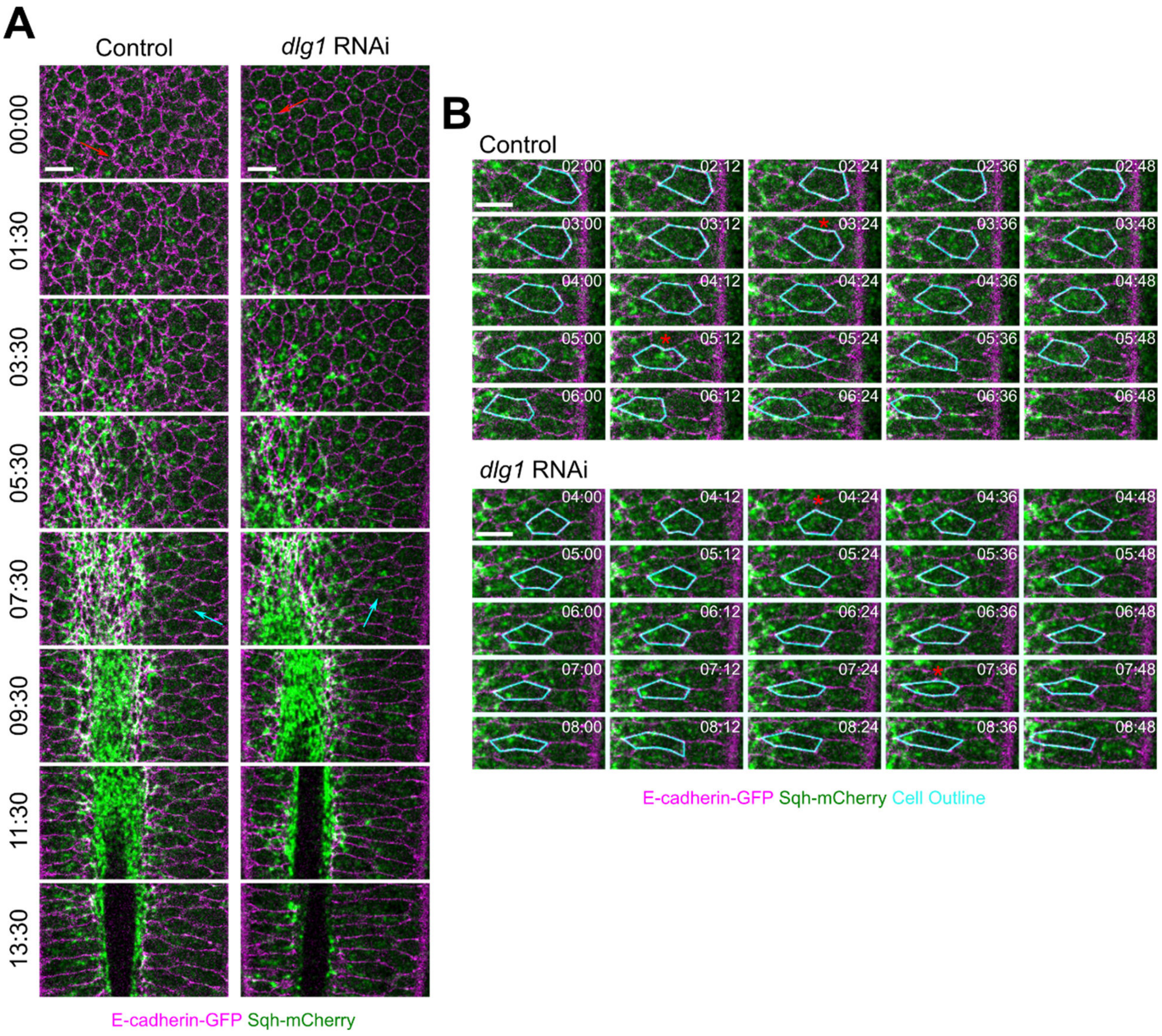


**Figure S5. Apical myosin accumulates at low levels in the flanking cells and exhibits cycles of coalescence and dispersion**

**(A)** Ventral surface view of a control embryo and a *dlg1* RNAi embryo expressing E-cadherin-GFP and Sqh-mCherry during ventral furrow formation. Red arrows point to apical myosin accumulation in constricting cells, while cyan arrows point to apical myosin accumulation in the flanking cells. Time zero is the onset of apical constriction. Scale bars: 10 μm. **(B)** Surface view of an individual flanking cell (cyan) from the control and *dlg1* RNAi embryo shown in A. Red asterisks indicate myosin pulses. Time zero is the onset of apical constriction. Scale bars: 10 μm.

### Figure S6

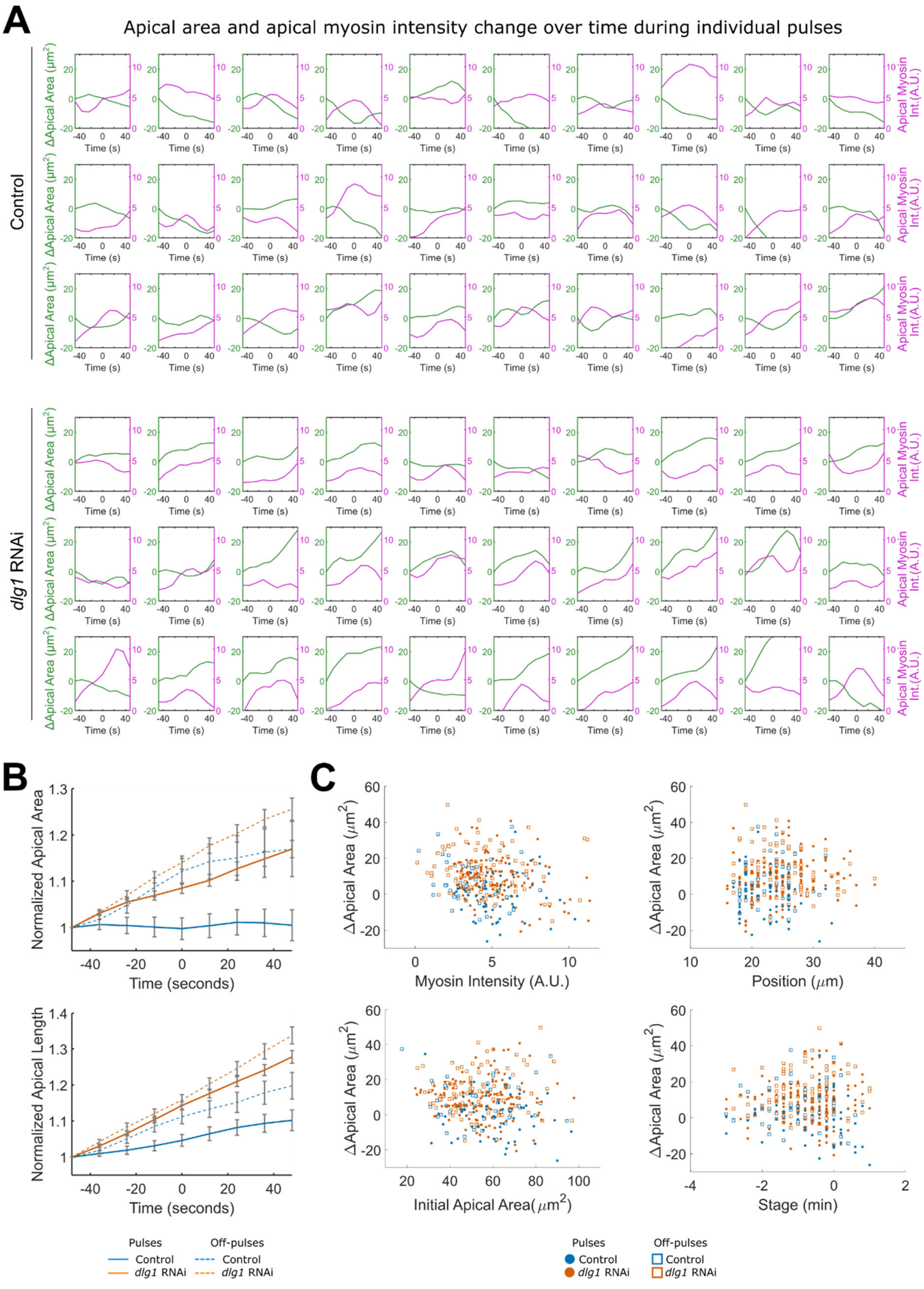
A
Apical area and apical myosin intensity change over time during individual pulses
Control
dlg1 RNAi
ΔApical Area (μm²)
Apical Myosin Int.(A.U.)
Time (s)
B
Normalized Apical Area
Time (seconds)
Normalized Apical Length
Pulses
Off-pulses
Control
dlg1 RNAi
C
ΔApical Area (μm²)
Myosin Intensity (A.U.)
Position (μm)
Initial Apical Area(μm²)
Stage (min)

**Figure S6. Apical myosin pulse behavior within the flanking cells**

**(A)** Example plots showing apical myosin intensity and apical cell area change over time for individual pulses. Time zero is the pulse peak. **(B)** Average trend of normalized apical area and length change over time for pulses and off-pulses in the flanking cells. 56 pulses and 41 off-pulses from 4 control embryos and 143 pulses and 100 off-pulses from 8 *dlg1* RNAi embryos were analyzed. **(C)** Distribution of apical area change for a pulse or off-pulse in the flanking cells as a function of i) myosin intensity at the pulse peak, ii) initial apical area of the cell, iii) position of the cell relative to the midline, and iv) the stage of the pulse relative to $T_{trans}$.

### Figure S7

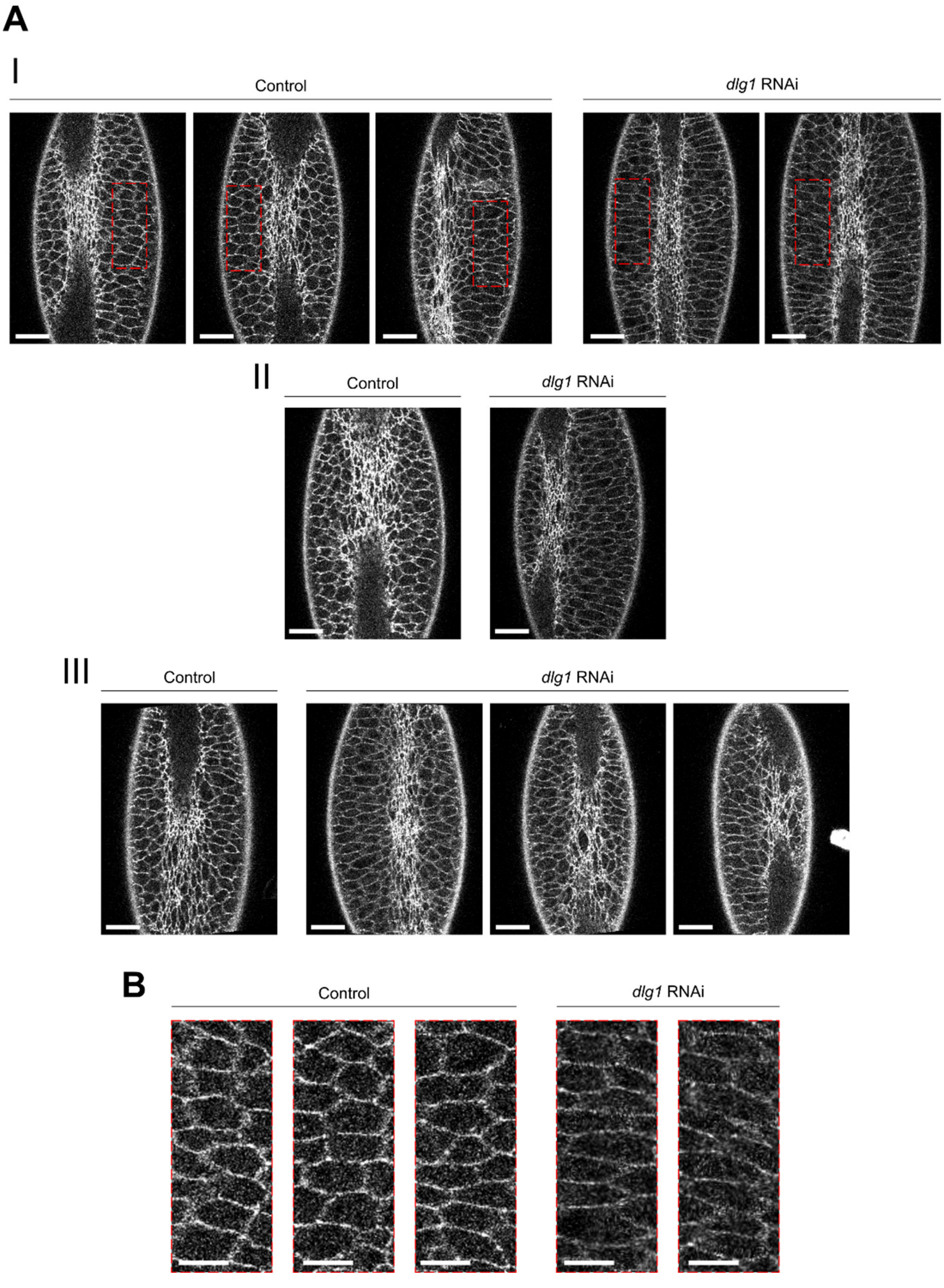
A
I
Control
dlg1 RNAi
II
Control
dlg1 RNAi
III
Control
dlg1 RNAi
B
Control
dlg1 RNAi

**Figure S7. E-cadherin levels are lower in the flanking cells of *dlg1* RNAi embryos compared to control embryos**

**(A)** Representative images of control and *dlg1* RNAi embryos expressing E-cadherin-GFP at $T_{trans}$ during ventral furrow formation taken on 3 different days (I, II, and III, respectively). Red box in I indicates position of flanking cells shown in B. Scale bars: 20 µm. **(B)** Zoomed in images of flanking cells in control and *dlg1* RNAi embryos marked by red boxes in A. Scale bars: 10 µm.

### Figure S8

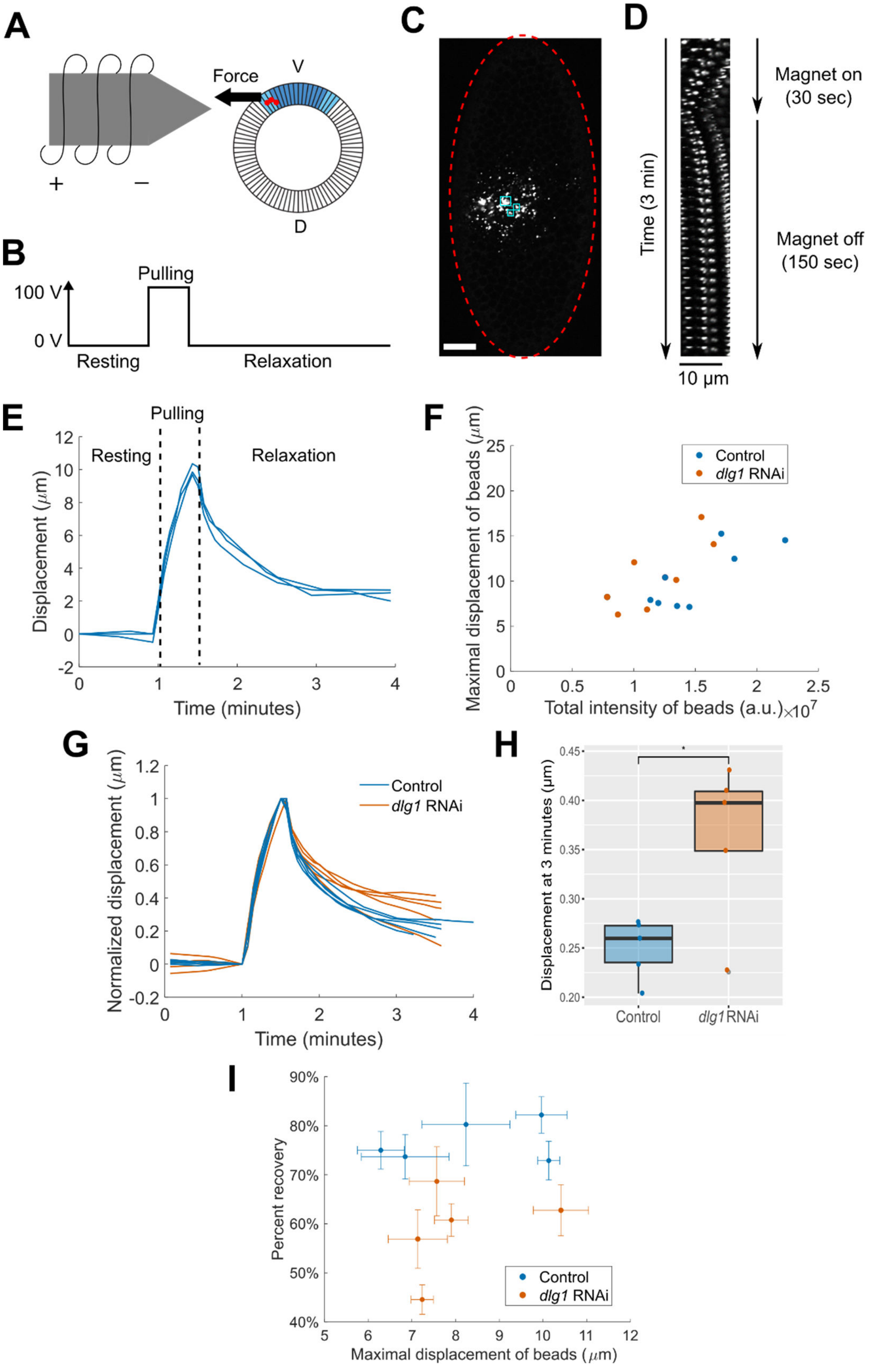
A
Force
V
D
+
−
B
Pulling
100 V
0 V
Resting
Relaxation
C
D
Time (3 min)
Magnet on
(30 sec)
Magnet off
(150 sec)
10 μm
E
Pulling
Resting
Relaxation
Displacement (μm)
Time (minutes)
F
Maximal displacement of beads (μm)
Total intensity of beads (a.u.) ×10^7
Control
dlg1 RNAi
G
Normalized displacement (μm)
Time (minutes)
Control
dlg1 RNAi
H
Displacement at 3 minutes (μm)
*
Control
dlg1 RNAi
I
Percent recovery
Maximal displacement of beads (μm)
Control
dlg1 RNAi

**Figure S8. The tissue of *dlg1* RNAi embryos is less elastic than that of control embryos as revealed by a magnetic tweezers-based approach**

**(A)** Schematic illustrating the magnetic tweezers experimental design. Cellularizing embryos were injected in the ventral-lateral region with magnetic beads that were 0.5 µm in diameter. An electromagnet was used to apply force and pull on the magnetic beads within the embryo. **(B)** The "On (100 Volts)" and "Off (0 Volts)" scheme of the magnetic tweezers during a typical experiment. **(C)** Example of a control embryo (outlined in red) injected with magnetic beads. Three clusters of beads (highlighted in cyan boxes) were tracked before, during, and after force application. Scale bar: 20 µm. **(D)** The kymograph for a cluster of magnetic beads illustrates the viscoelastic response of the tissue to a pulling force. **(E)** The displacement over time curves for the beads in the embryo marked in panel C. Each curve corresponds to one tracked bead cluster. **(F)** Total bead intensity for individual control ($n = 8$) and *dlg1* RNAi embryos ($n = 8$) was measured and plotted against the maximal displacement of the tracked bead clusters for each experiment. Maximal magnetic bead displacement positively correlates with total bead intensity. **(G)** Normalized average displacement curves for individual control ($n = 5$) and *dlg1* RNAi embryos ($n = 5$), where the maximal displacement is within a 6 – 11 µm range. For each embryo, three clusters of beads near the center of the bead distribution were tracked. **(H)** Boxplots showing the displacement of the beads at 3 minutes for individual control ($n = 5$) and *dlg1* RNAi embryos ($n = 5$) shown in panel F. Student t-test, unpaired, one-tail. *: $p < 0.05$; **: $p < 0.01$; ***: $p < 0.005$. **(I)** Maximal magnetic bead displacement plotted against percent recovery for control ($n = 5$) and *dlg1* RNAi embryos ($n = 5$) shown in panel F. Each data point corresponds to the average measurement in one embryo. Error bar corresponds to the standard deviation of the three bead clusters tracked in each embryo.

#### Figure S9

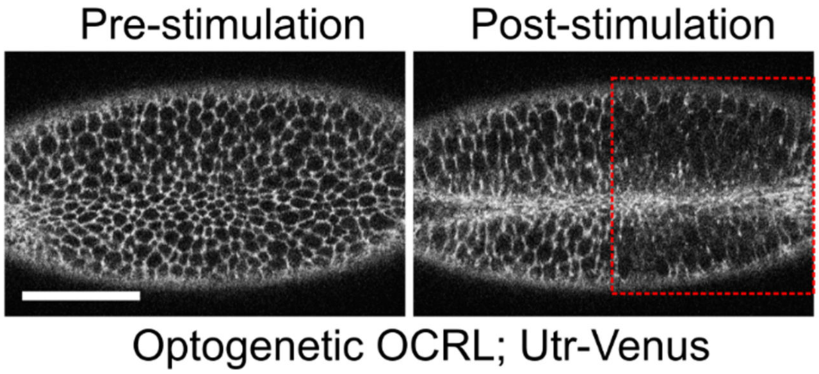


**Figure S9. Multiphoton stimulation in embryos expressing the optogenetic OCRL components results in down regulation of F-actin**

Two photon stimulation of an embryo expressing Utr-Venus and the optogenetic OCRL components (CIBNpm without GFP, CRY2-mCherry-OCRL) results in a rapid loss of Utr-Venus in the stimulated region (red box).

#### Supplemental movies

**Movie 1: Depletion of Dlg1 delays the transition from apical constriction to invagination (surface view).**

Wild type embryo (left) and *dlg1* RNAi embryo (right) undergo apical constriction at a similar rate. However, the transition to invagination is delayed in the *dlg1* RNAi embryo. Note that the apical domain of the flanking, non-constricting cells adjacent to the constriction domain are over-stretched in the *dlg1* RNAi embryo. Cell membranes are labeled with E-cadherin-GFP. T = 00:00 (mm:ss) marks the onset of gastrulation.

**Movie 2: Depletion of Dlg1 delays the transition from apical constriction to invagination (cross-section view).**

A *dlg1* RNAi embryo (right) shows delayed transition from apical constriction to invagination. A wildtype embryo is shown on the left for comparison. The cross-section views were generated by re-slicing the 3D image stacks acquired over time. Cell membranes are labeled with E-cadherin-GFP. T = 00:00 (mm:ss) marks the onset of gastrulation.

**Movie 3: The apical myosin network behaves normally in the constricting cells of *dlg1* RNAi embryos.**

Confocal images of a representative control and *dlg1* RNAi embryo expressing Sqh-mCherry (magenta) and E-cadherin-GFP (green). Maximum projections of the *en face* view for Sqh-mCherry and a single apical plane for E-cadherin-GFP are shown. Apical myosin accumulation and the spatial organization of the myosin network are comparable between the control and the *dlg1* RNAi embryos during apical constriction. T = 0:00 (mm:ss) corresponds to the onset of gastrulation.

**Movie 4: Knockdown of Dlg1 affects apical myosin contractions in the flanking cells.**

Ventral surface view of a control embryo and a *dlg1* RNAi embryo expressing E-cadherin-GFP (magenta) and Sqh-mCherry (green) during late apical constriction. For each movie, two representative flanking cells are highlighted in magenta. T = 0:00 (mm:ss) corresponds to the onset of gastrulation.

**Movie 5: Laser ablation of the flanking, non-constricting cells delays the transition from apical constriction to invagination.**

Laser ablation of two strips of flanking cells adjacent to the constriction domain in wildtype embryos during apical constriction. A stage matched, non-ablated embryo is shown on the left for comparison. Top: cross-section view; Bottom: ventral surface view. Cell membranes are visualized using E-cadherin-GFP. T = 00:00 (mm:ss) corresponds to the transition from apical constriction to invagination in control embryos.

**Movie 6: Optogenetic disruption of F-actin in the flanking, non-constricting cells delays the transition from apical constriction to invagination.**

An embryo expressing the CIBN-pm and CRY2-mCherry-OCRL optogenetic constructs and the myosin marker, Sqh-mCherry, was stimulated on one side (adjacent to the constriction domain) during an early stage of apical constriction (right panel). A stage-matched, unstimulated embryo is shown on the left for comparison. Top: cross-section view; Bottom: ventral surface view. T =

00:00 (mm:ss) corresponds to the transition from apical constriction to invagination in control embryos.
